# Comparative genomics of peroxisome biogenesis proteins: making sense of the PEX mess

**DOI:** 10.1101/2020.12.16.423121

**Authors:** Renate L.M. Jansen, Carlos Santana Molina, Marco van den Noort, Damien P. Devos, Ida J. van der Klei

## Abstract

PEX genes encode proteins involved in peroxisome biogenesis and proliferation. Using a comparative genomics approach, we clarify the evolutionary relationships between the 37 known PEX proteins in a representative set of eukaryotes, including all common model organisms, pathogenic unicellular eukaryotes and human. A large number of previously unknown PEX orthologs were identified. We analysed all PEX proteins, their conservation and domain architecture and defined the minimum set of PEX proteins that is required to make a peroxisome. The molecular processes in peroxisome biogenesis in different organisms were put into context, showing that peroxisomes are not static organelles in eukaryotic evolution. Organisms that lack peroxisomes still contain a few PEX proteins, which probably play a role in alternative processes. Finally, the relationships between PEX proteins of two large families, the Pex11 and Pex23 families, were clarified, thereby contributing to the understanding of their complicated and sometimes incorrect nomenclature. We provide an exhaustive overview of this important eukaryotic organelle.

## Introduction

Peroxisomes occur in almost all eukaryotes. Their number, size and protein composition are highly variable. In lower eukaryotes, such as yeast, peroxisome proliferation is stimulated by specific growth substrates. In higher eukaryotes, peroxisome abundance and composition vary with organism, tissue and developmental stage. Conserved peroxisomal pathways are the β-oxidation of fatty acids and hydrogen peroxide degradation. Examples of specialized pathways are the biosynthesis of bile acids and ether lipids in man, photorespiration in plants and the biosynthesis of antibiotics in certain filamentous fungi (Smith & Aitchison, 2013). The crucial role of peroxisomes for human health is illustrated by the occurrence of inborn errors that cause severe diseases and are often lethal. However, roles in non-metabolic processes such as ageing, anti-viral defence and cancer show that the significance of peroxisomes in human health goes far beyond the relatively rare inherited peroxisomal disorders (Islinger et al, 2018).

Peroxisomes are very simple organelles that consist of a protein rich matrix surrounded by a single membrane. Peroxisomal enzymes almost exclusively occur in the matrix. The membrane contains transporters, pores for solute transport and proteins involved in diverse processes such as matrix and membrane protein sorting, organelle fission and movement (*figure 1*).

**Figure 1.**
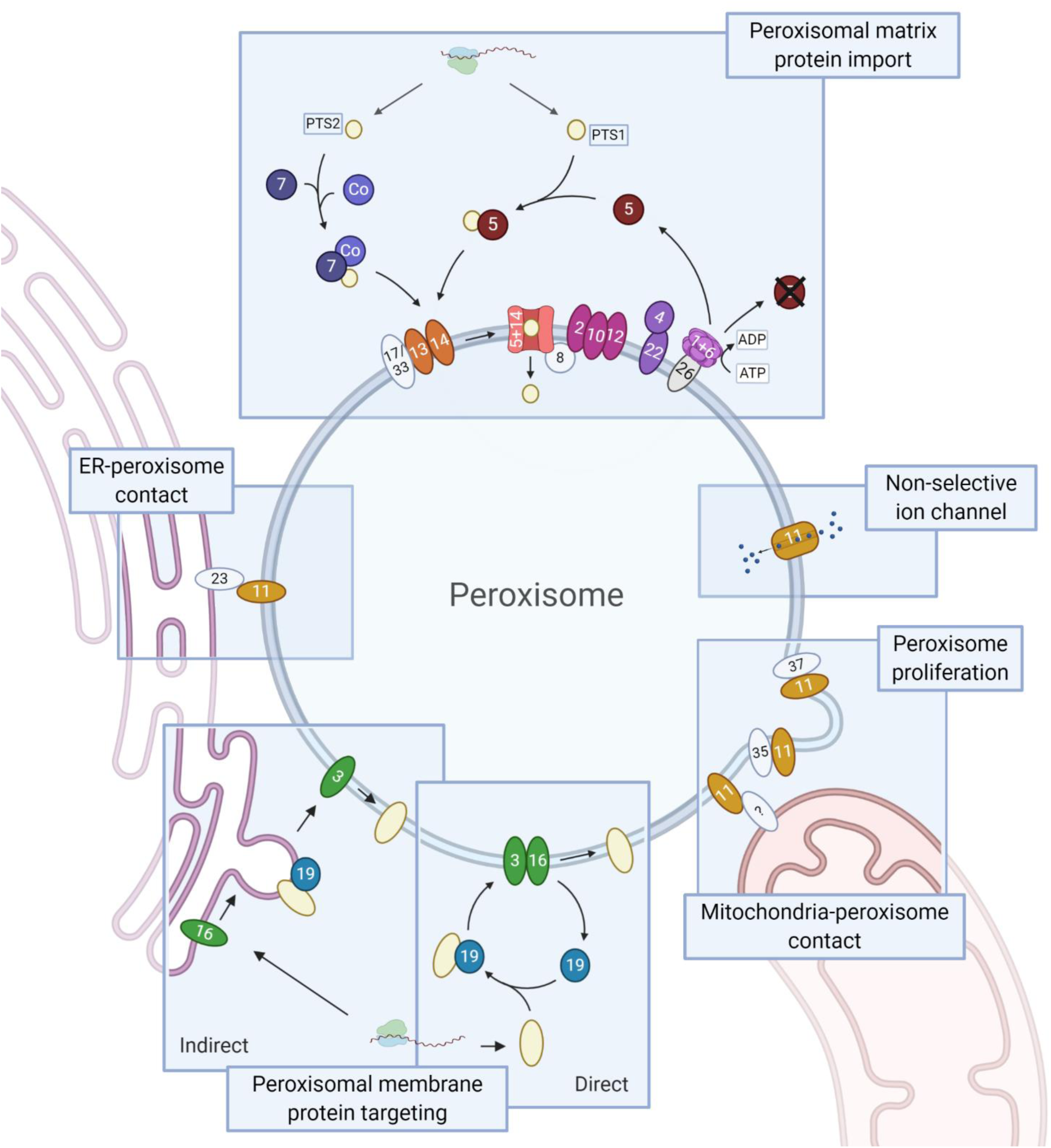
Schematic representation of the PEX proteins. Core conserved PEX proteins (shapes in dark colours, names in white), fungi-specific proteins (light, names in black) and the moderately conserved PEX protein Pex26/15 (grey, name in black, which is only present in metazoa and fungi) are depicted. Membrane proteins are ovals, soluble proteins round. **Matrix protein import.** Peroxisomal matrix proteins contain a peroxisomal targeting signal (PTS) that is recognized by cytosolic receptors: a C-terminal PTS1 or (less commonly) an N-terminal PTS2, recognized by PEX5 and PEX7 respectively. PTS2 import involves a co-receptor (Co): PEX5 (animals, plants and protists), PEX18/21 (*S. cerevisiae*) or PEX20 (fungi). Next, the receptor-cargo complex associates with the docking complex, consisting of PEX13/14 (and in fungi PEX17 or PEX33). Upon cargo translocation and release, the PTS (co-)receptor is ubiquitinated and recycled. Ubiquitination involves the ubiquitin conjugating enzyme (E2) PEX4 (recruited to the membrane by PEX22) and the ubiquitin ligase (E3) activities of the RING finger complex, consisting of PEX2/10/12. Receptor extraction requires the AAA+ ATPase complex PEX1/6, which is recruited to the membrane via PEX26 (PEX15 in *S. cerevisiae*). PEX8 bridges the docking and RING finger complexes, and functions in receptor-cargo dissociation. **Peroxisomal membrane protein (PMP) sorting** involves PEX3, PEX19 and PEX16. PMPs can sort directly to peroxisomes or indirectly via the ER. In the direct pathway PEX19 acts as receptor/chaperone, while it functions at the ER in PMP sorting via the indirect pathway. The **Pex11 protein family** (all show as PEX11) and the fungal peroxins PEX35 and PEX37 have been mainly implicated in peroxisome proliferation. Pex11 family proteins are also present in mitochondria-peroxisome contact sites and PEX11 functions as non-selective ion channel. Members of the fungal **Pex23 protein family** localize to the ER and are involved in the formation of peroxisome-ER membrane contact sites. Created with BioRender.com.

In 1996, the term peroxin was coined for proteins “involved in peroxisome biogenesis (inclusive of peroxisomal matrix protein import, membrane biogenesis, peroxisome proliferation, and peroxisome inheritance)” (Distel et al, 1996). Peroxins are encoded by PEX genes and also called PEX proteins. So far, 37 PEX proteins have been described. Some are highly conserved, whereas others only occur in a limited number of species. Since 1996, tremendous progress has been made in our understanding of the molecular mechanisms involved in peroxisome biology. However, with the increasing number of PEX proteins, their nomenclature became more and more complex (Smith & Aitchison, 2013).

Here, we present an exhaustive up-to-date overview of all the PEX protein families. We analysed PEX proteins in a highly diverse set of eukaryotes, including all common model organisms, pathogenic unicellular eukaryotes and higher eukaryotes. Using this information, we combine phylogenetic reconstructions with other protein features (e.g., Pfam domain, protein disorder and transmembrane domain predictions) to understand the evolution of these proteins, clarifying certain inconsistencies in the nomenclature of PEX proteins. Important questions that we answer are (i) how are the different PEX genes conserved across eukaryotes, (ii) what is the minimum set of PEX genes to make a canonical peroxisome and (iii) what are the typical features of the PEX proteins.

## Results

The proteomes of 38 eukaryotes were investigated to identify all PEX proteins known to date. Not all eukaryotes contain peroxisomes (Žárský & Tachezy, 2015) and several protist species of our initial analysis were found to lack most PEX proteins, namely *Cryptosporidium parvum, Theileria annulata, Babesia bovis, Monosiga brevicollis, Plasmodium falciparum, Blastocystis hominis* and *Entamoeba histolytica*. To facilitate comparison between species containing and (likely) lacking peroxisomes, the latter species was included in further analyses, but all others likely lacking peroxisomes were omitted. *Table 1* shows the 31 remaining species containing peroxisomes, plus *Entamoeba histolytica*. An overview of all orthologs identified can be found in *table S1*.

**Table 1:**
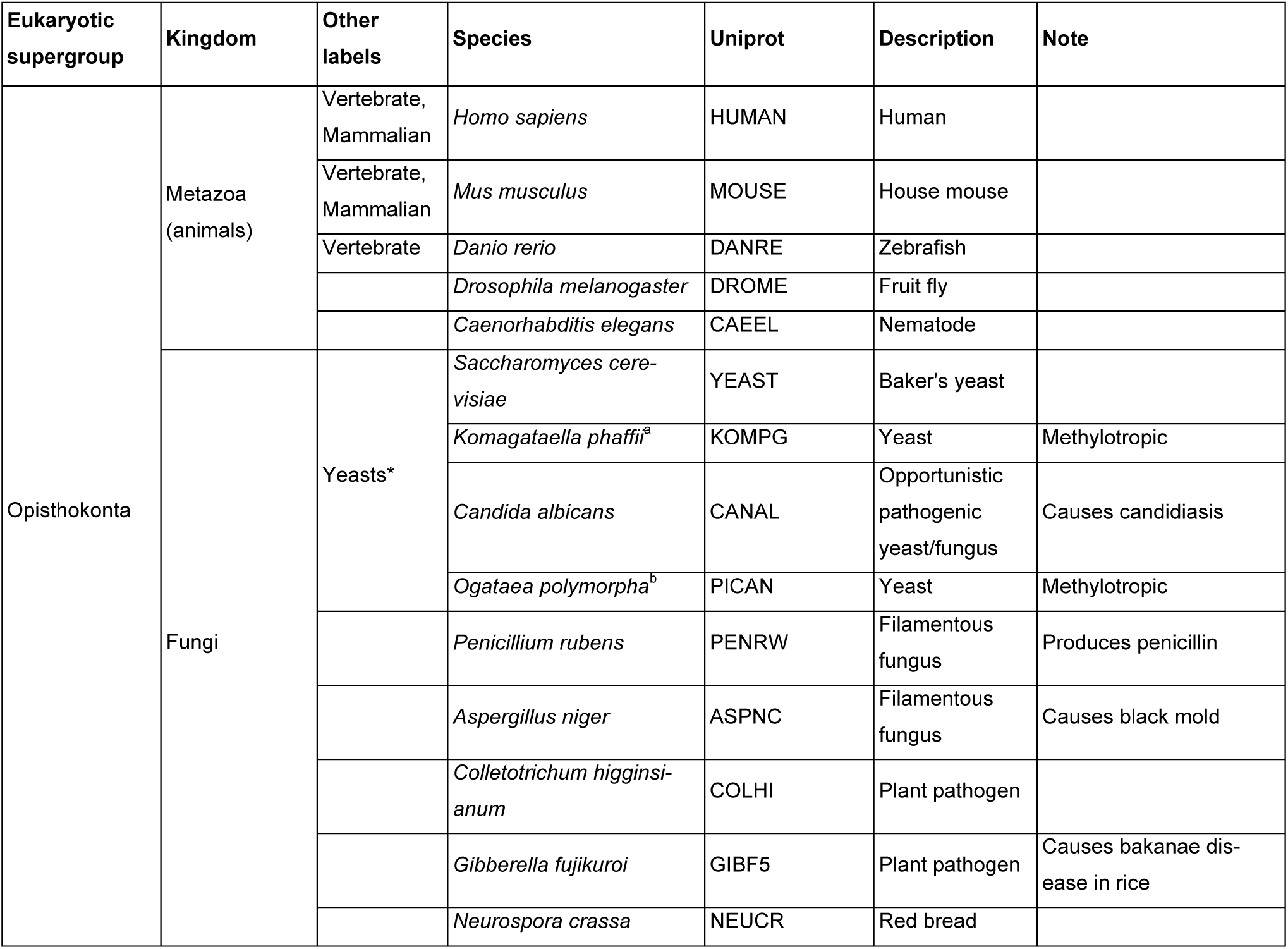
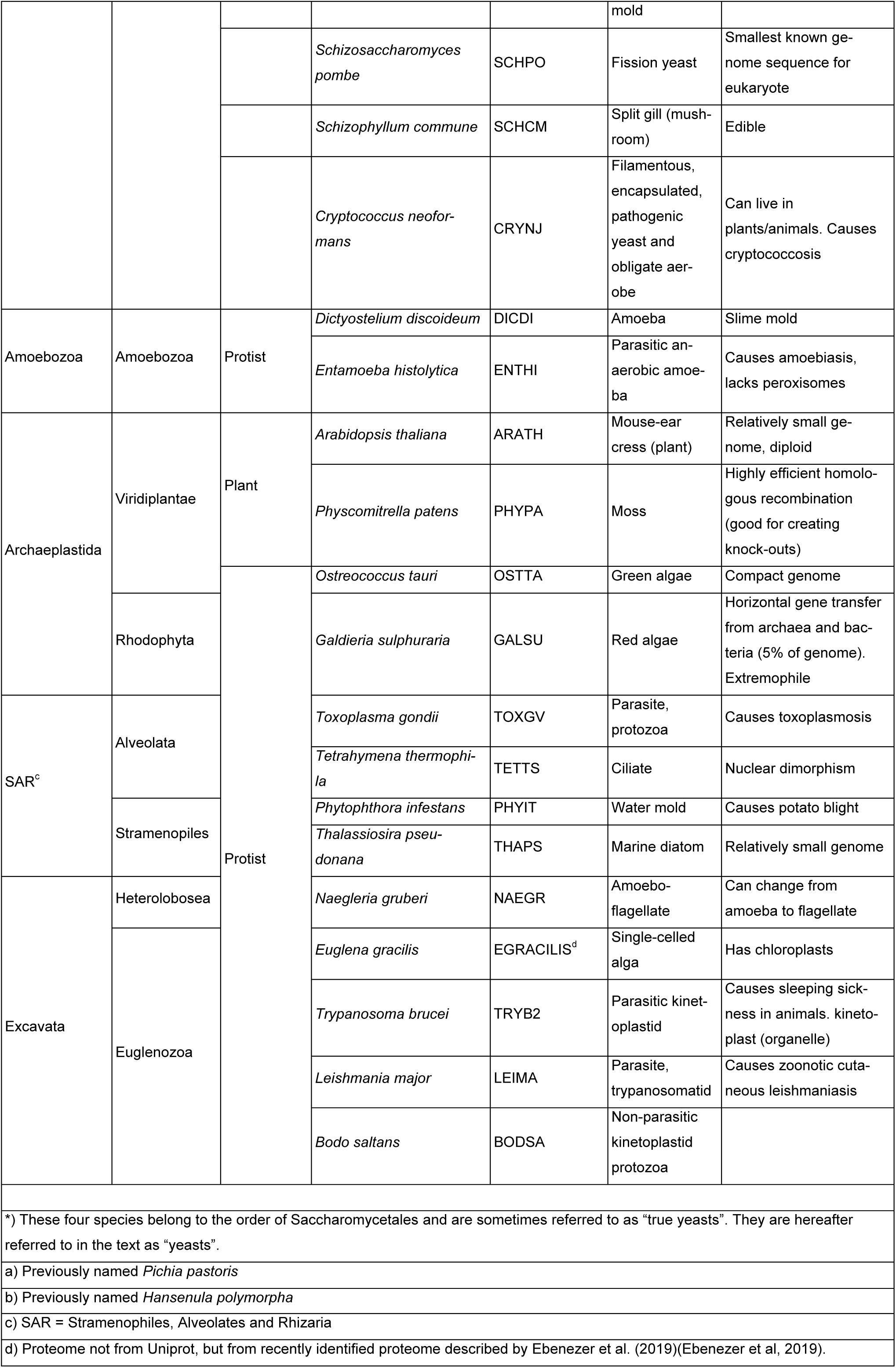
Overview of proteomes investigated.

| Eukaryotic supergroup | Kingdom | Other labels | Species | Uniprot | Description | Note |
| --- | --- | --- | --- | --- | --- | --- |
| Opisthokonta | Metazoa (animals) | Vertebrate, Mammalian | *Homo sapiens* | HUMAN | Human |  |
|  |  | Vertebrate, Mammalian | *Mus musculus* | MOUSE | House mouse |  |
|  |  | Vertebrate | *Danio rerio* | DANRE | Zebrafish |  |
|  |  |  | *Drosophila melanogaster* | DROME | Fruit fly |  |
|  |  |  | *Caenorhabditis elegans* | CAEEL | Nematode |  |
|  | Fungi | Yeasts* | *Saccharomyces cerevisiae* | YEAST | Baker's yeast |  |
|  |  |  | *Komagataella phaffii*<sup>a</sup> | KOMPG | Yeast | Methylotrophic |
|  |  |  | *Candida albicans* | CANAL | Opportunistic pathogenic yeast/fungus | Causes candidiasis |
|  |  |  | *Ogataea polymorpha*<sup>b</sup> | PICAN | Yeast | Methylotrophic |
|  |  |  | *Penicillium rubens* | PENRW | Filamentous fungus | Produces penicillin |
|  |  |  | *Aspergillus niger* | ASPNC | Filamentous fungus | Causes black mold |
|  |  |  | *Colletotrichum higginsianum* | COLHI | Plant pathogen |  |
|  |  |  | *Gibberella fujikuroi* | GIBF5 | Plant pathogen | Causes bakanae disease in rice |
|  |  |  | *Neurospora crassa* | NEUCR | Red bread |  |

|  |
| --- |
| c) SAR = Stramenophiles, Alveolates and Rhizaria |
| d) Proteome not from Uniprot, but from recently identified proteome described by Ebenezer et al. (2019)(Ebenezer et al, 2019). |

### 1. Distribution and general description of PEX proteins across eukaryotic lineages

The results of our computational survey are summarized in *figure 2*. We detect a core of PEX proteins that are **broadly conserved** in all eukaryotic lineages, encompassing PEX3/19/16 (peroxisomal membrane protein (PMP) sorting), PEX1/6, PEX2/10/12, PEX13/14 and PEX5/7 (matrix protein import) and proteins of the Pex11 family (peroxisome proliferation and contact sites). Some detected **absences** are probably real. On the other hand, in other cases the function of missing PEX proteins may be taken over by other homologous proteins. For instance, the function of the ubiquitin conjugating enzyme (E2 enzyme) PEX4 in receptor ubiquitination is performed by proteins of the E2D family in Metazoa, which lack a PEX4 ortholog (Grou et al, 2008). Similarly, the function of PEX26 is complemented by the homologous protein APEM9 in plants (Cross et al, 2016) and PEX15 in *S. cerevisiae.* Furthermore, we observe an important bias towards fungi (yeasts and filamentous fungi) reflected in the large number of PEX proteins that are specific to fungi, such as PEX8, PEX20/18/21 and the Pex23 family (*figure 2*). This is a result of the fact that the large majority of studies investigating peroxisomes, in particular their biogenesis, have been performed in yeast models such as *S. cerevisiae, O. polymorpha, K. phaffii* and *Yarrowia lipolytica*.

**Figure 2:**
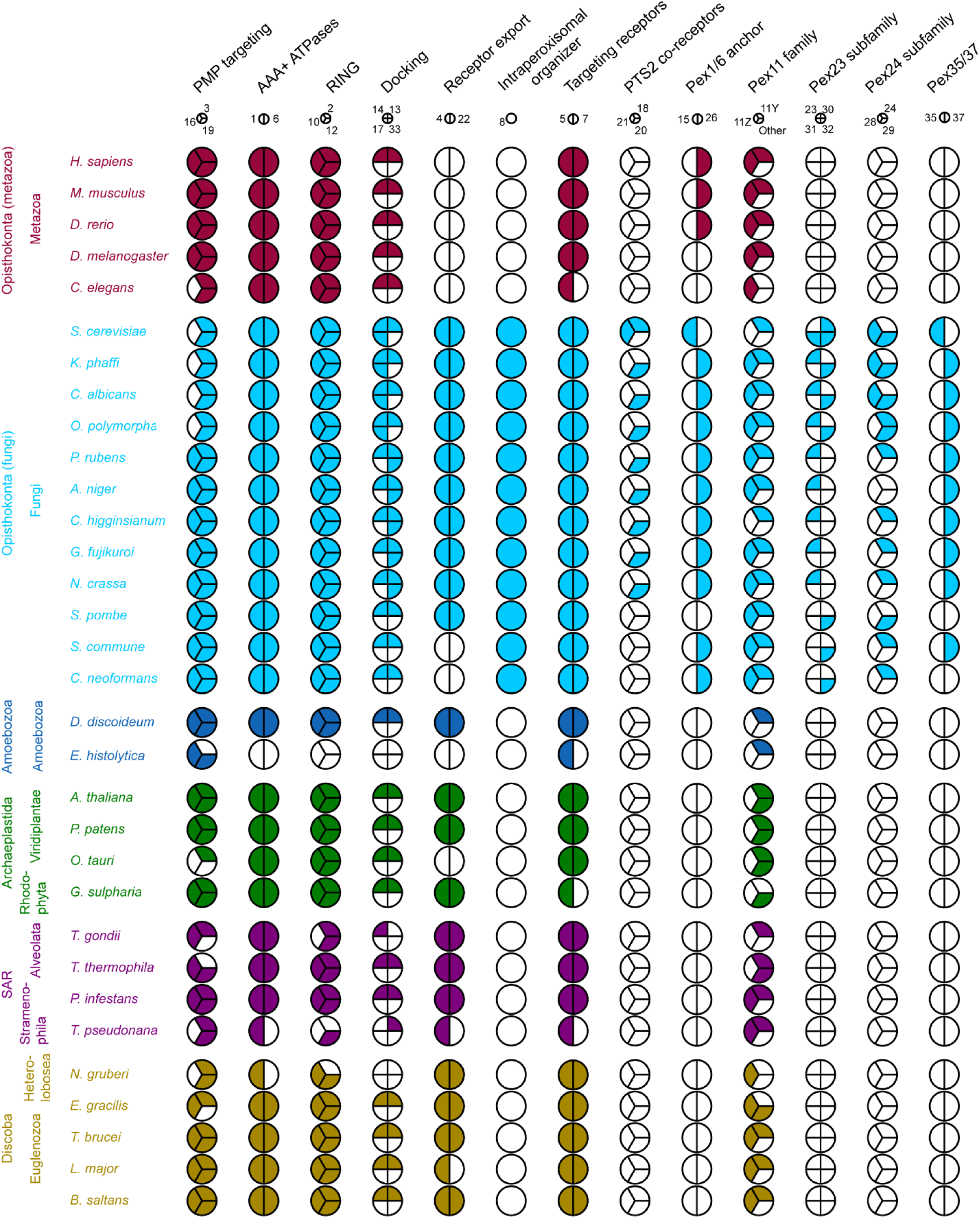
Coulson plot demonstrating the presence (filled) or absence (empty) of PEX protein orthologs in 32 eukaryotic proteomes. PEX proteins are divided into functional groups (columns) including homologous and non-homologous proteins, represented by a pie. Every wedge represents a PEX protein, with the exception of the Pex11 family, where each wedge represents a subfamily. The PEX11Y subfamily contains among others fungal PEX11/25/27/34/36 and mammalian PEX11α/β. The PEXZ subfamily contains fungal PEX11C and mammalian PEX11γ. Pex11 family proteins that do not belong to the PEX11Y or PEX11Z subfamilies are placed in “Other”. Organisms are grouped by eukaryotic supergroup (colour-coded for clarity) and kingdom. PEX proteins are designated by their number.

We analysed the structural features of the PEX proteins (see *table 2*). Structural protein disorder seems to be a common feature among some PEX proteins. In some of them, structural disorder is only predicted for a short fragment, but others like PEX19, PEX18/20/21, PEX14/33-13 are predicted as almost entirely disordered. Also, transmembrane helical domains are usually present in certain PEX proteins, such as PEX3, PEX14 and PEX26. Several PEX proteins have common eukaryotic structural domains, like the E2 enzyme PEX4 and the AAA+ ATPase domain present in PEX1 and PEX6. We also detect several functional domain associations, such as the RING finger (zinc finger) domain in PEX2/10/12 and SH3 domains in PEX13, both being involved in signal transduction and controlling protein-protein interactions. Other recognizable fold types in PEX proteins include α-solenoid formed by the TPR repeat domains in PEX5 and the β-propeller formed by WD40 repeats in PEX7.

**Table 2:** Overview of PEX proteins and their main features.

| Main members | Actual groups | Naming inconsistency | Protein disorder | Trans-membrane | Pfam | Distribution |
| --- | --- | --- | --- | --- | --- | --- |
| PEX1, 6 | PEX1, 6 |  | Some regions | - | AAA, AAA_lid_3, PEX-1N | Eukaryotes |
| PEX2, 10, 12 | PEX2, 10, 12 |  | Some regions | + | Pex2_Pex12, Zn-finger | Eukaryotes |
| PEX3 | PEX3 |  | Some regions | + | peroxin-3 | Eukaryotes |
| PEX4 | PEX4 |  | - | - | UB_con | Eukaryotes |
| PEX5, 9 | PEX5 | PEX9 is actually a duplication of PEX5, specific to *S. cerevisiae* | +, N-terminal | - | TPR | Eukaryotes |
| PEX7 | PEX7 |  | - | - | WD40 (B-propeller) | Eukaryotes |
| PEX8 | PEX8 |  | - | - | unknown | Fungi |
| Pex11 family, including PEX25, 27, 34, 36 | Pex11Y subfamily (incl. group PEX25/27/34/36), 11Z subfamily, Other | Current PEX11A/B names in different organisms are inconsistent. PEX25, 27, 34 and 36 belong to the same fungi-specific paralog group | - | + | PEX11 | Pex11Y-Z Eukaryotes / PEX25/27/34/36 Fungi |
| PEX13 | PEX13 |  | +, (N-terminal) | + | Pex13, SH3 | Eukaryotes |
| PEX14, 33 | PEX14, 33 |  | + | + | Pex14_N | PEX14-Eukaryotes / PEX33-fungi |
| PEX15, 26 | PEX15, 26 |  | Some regions | + | Pex26 | Fungi-metazoa |
| PEX16 | PEX16 |  | Some regions | + | Pex16 | Eukaryotes |
| PEX17 | PEX17 |  | - | + | unknown | Fungi |
| PEX19 | PEX19 |  | + | +? | Pex19 | Eukaryotes |
| PEX18, 20, 21 | PEX20 | PEX18 and PEX21 of *S. cerevisiae* are the result of the duplication of the fungi specific PEX20 form | + | - | unknown | Fungi |
| PEX22 | PEX22 |  | - | + | peroxin_22 | Fungi-Plants-Protist |
| PEX23, 24, 28, 29, 30, 31, 32, 23-like, TECPR1 | TCPR1, Pex23 subfamily, Pex24 subfamily, 23-like (sporulation) | PEX28 and PEX24 are actually the same protein in different organisms (*O. polymorpha* and *S. cerevisiae*). PEX30 and PEX31 are a specific duplication in *S. cerevisiae* of PEX23 form | Some regions | +, (TCPR1 does not) | pex24p, Hyd_WA (TCPR1) | Fungi / TECPR1 metazoa |
| PEX35 | PEX35 |  | - | + | unknown | *Saccharomycetaceae* |
| PEX37 | PEX37 |  | Some, C-terminal | + | Mpv17_PMP22 | Fungi |

The actual groups indicate to the main groups (deep paralogs) identified in phylogenetic reconstructions. Protein disorder was predicted using IUPRED and transmembrane helices through TMHMM software. Functional protein domains
annotated using Pfam database. Question marks indicate those features that were present in a subset of protein sequences from our data set.

The functional diversification of proteins is caused by the duplication of the respective genes. This is one of the main sources of cellular complexity and development. This process is called paralogization, where paralogous proteins are those having a common origin, i.e., belonging to the same protein family. These gene duplications (paralogizations) can be ancestral (deep paralogs) or they can be asynchronous during evolution: appearing later and being restricted to specific taxonomic clades (in-paralogs). The paralogization of PEX proteins seems to have been relevant for the development of peroxisomes in Eukarya domain. Indeed, some of these paralogizations preceded the diversification of eukaryotes, like the peroxins of the AAA+ ATPase protein family PEX1/6, the RING finger proteins PEX2/10/12 and proteins of the Pex11 protein family. On the other hand, some other PEX proteins have been duplicated in specific eukaryotic taxons. These proteins have often been inconsistently named, since newly discovered proteins were sometimes given a new number. This should be kept in mind when studying such proteins. For instance, the *S. cerevisiae* PEX9 is actually a copy of PEX5 (in-paralogs, not ancestral duplication in fungi). Similarly, the fungal PTS2 co-receptors PEX18/20/21 should be considered as a single group: PEX18 and PEX21 of *S. cerevisiae* are actually the result of a duplication of the ancestral PEX20 form. The PEX23 family is specifically found in fungi and encompasses multiple copies in specific organisms, such as PEX30/31/32 and PEX28/29 in *S. cerevisiae*, resulting from the duplication of PEX23 and PEX24, respectively. In the previous examples, different proteins derived from the same ancestral protein, i.e., belonging to the same protein family, have received different numbers. On the other hand, the opposite has happened for certain other PEX proteins. Many members of the Pex11 family have the same number, but were given a different appendix instead: for instance, PEX11α/β/γ or PEX11A/B/C/D/E. We detected that these paralogs originated from independent paralogizations in different lineages, but their naming does not always reflect this. For instance, fungal PEX11C belongs to the same subfamily as human PEX11γ, but PEX11C from *A. thaliana* does not. Similarly, *A. thaliana* PEX11A is not equivalent to human PEX11α. Based on phylogenetic reconstructions, we propose that two different subfamilies can be distinguished within the Pex11 family. In addition, the Pex11 family includes an in-paralog group specific to fungi, containing PEX25/27/34/36.

Therefore, in some cases, the nomenclature ascribed to the PEX protein paralogizations could lead to confusion, because there is no uniformity in the way in which paralogous, in-paralogous or non-paralogous/unrelated proteins have been named. Furthermore, some paralogizations have led to paralogs of PEX proteins that may no longer function in peroxisome biology. For instance, vertebrates express a PEX5 paralog called PEX5R (TRIP8b), whose only known function is the regulation of hyperpolarization-activated cyclic nucleotide-gated (HCN) channels - key modulators of neuronal activity(Han et al, 2020).

Taking into account all of the above, we review the role of these PEX proteins below, in order to gain a comprehensive understanding of their functional classification.

### 2. A core set of PEX proteins is broadly conserved in Eukaryotes

A core set of PEX proteins is broadly conserved across all eukaryotic lineages, encompassing proteins involved in PMP sorting (PEX3, PEX19 and PEX16), matrix protein receptors (PEX5 and PEX7), components of the receptor docking site (PEX13 and PEX14), enzymes involved in receptor ubiquitinylation (PEX2, PEX10, PEX12 and PEX4), two AAA-ATPases that play a role in receptor recycling (PEX1 and PEX6) and a protein family involved in peroxisome proliferation (Pex11 family). The function of these conserved PEX proteins is central to peroxisome biology and thus maintained. In the following section, we will review how these processes define the biology of the canonical peroxisomes as well as the mechanistic models proposed in the field. Furthermore, we describe variations in the repertoire of PEX proteins in certain eukaryotes.

#### Sorting of PMPs (PEX3, PEX19 and PEX16)

Only three PEX proteins (PEX3, PEX16 and PEX19) are known to be involved in targeting of PMPs. Two mechanisms of PMP sorting to the peroxisome membrane have been described (see *figure 1*; for a detailed review, see (Jansen & van der Klei, 2019)). According to the direct sorting model, PEX19 binds to newly translated PMPs in the cytosol. In this pathway PEX19 acts as a chaperone and cycling receptor (Jansen & van der Klei, 2019). The PEX19-PMP complex binds to the PMP PEX3 and is subsequently inserted in the membrane by a currently unknown mechanism. In the indirect pathway, PMPs traffic first to the ER and accumulate at a subdomain, where PMP containing vesicles bud off. PEX3 plays a role in the intra-ER sorting of PMPs (Fakieh et al, 2013), while PEX19 is important for vesicle budding (Agrawal et al, 2016, Van Der Zand et al, 2012). PEX16 plays a role in the indirect pathway (Hua & Kim, 2016). Notably, PEX3 is also involved in a host of other functions, including pexophagy, peroxisome retention during yeast budding and the formation of contacts between peroxisomes and vacuoles. In all these processes, PEX3 recruits proteins to the peroxisomal membrane (e.g. Atg30/36, Inp1) (Jansen & van der Klei, 2019).

Our computational survey shows that PEX3, PEX19 and PEX16 are conserved well, with a few exceptions, suggesting minor variations in mechanisms of PMP sorting. For instance, PEX16 is widely conserved, but is absent in all (investigated) yeast species, *C. elegans* and several protists. A characteristic motif in PEX19 orthologs of many species is a **CaaX** box at the C-terminus. Farnesylation of this motif causes conformational changes in PEX19 and increases its binding affinity for PMPs (Emmanouilidis et al, 2017, Rucktaschel et al, 2009). Previous studies in *S. cerevisiae* and humans are contradictory regarding the importance of this post-translational modification for peroxisome function (Rucktaschel et al, 2009, Schrul & Kopito, 2016, Vastiau et al, 2006). Interestingly, Schrul & Kopito (2016) found that the CaaX box of human PEX19 was important for targeting of lipid droplet protein UBXD8, but not for peroxisome biogenesis (Schrul & Kopito, 2016). We checked if the CaaX box is present in all eukaryotes. We found that while this motif is present in all animals, plants and fungi, it is absent (or difficult to align) in many protists, like euglenozoa and amoebozoa, despite these organisms expressing the enzyme required for farnesylation (see e.g. (Buckner et al, 2002)) (*figure 3*). Interestingly, putative PEX19 orthologs were also identified in *Entamoeba histolytica* and *M. brevicollis*, despite these species very likely lacking peroxisomes. This may suggest an alternative function for PEX19, unrelated to peroxisomes.

**Figure 3.**
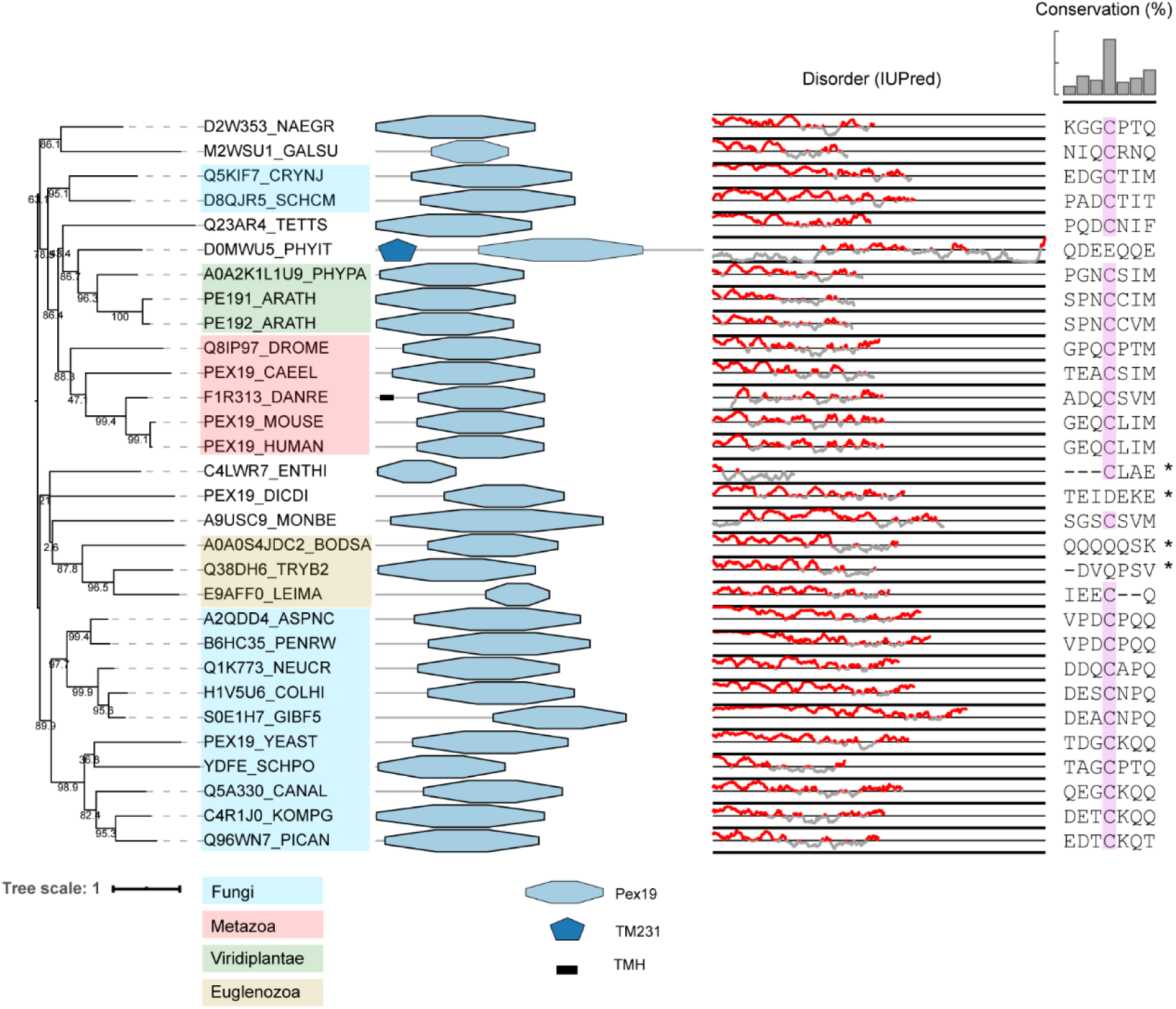
Phylogeny and protein features of PEX19 orthologs. The phylogeny is rooted at mid-point to ease the visualization and labels of the main taxonomic groups are coloured according to the legend. Note that the topology does not necessarily reflect the actual evolutionary trajectory of such proteins. Protein domain architecture is defined by pfam annotations and transmembrane helices (TMH) according to TMHMM software. The line-dot plot, indicates the regions predicted to be disordered (red) and not disordered (grey). The sequence alignment shows the conservation of the CaaX box in PEX19 orthologs of distant eukaryotes, with ‘C’ denoting Cys, ‘a’ an aliphatic residue and ‘X’ usually being a Ser, Thr, Gln, Ala or Met. Asterisk indicates forced alignments manually.

#### Matrix protein receptors (PEX5 and PEX7)

Newly synthesized matrix proteins are first recognized by their cytosolic **peroxisomal targeting signal (PTS) receptor**. The majority of peroxisomal matrix proteins contain a PTS1 or a PTS2, recognized by PEX5 and PEX7 respectively.

PEX7 contains WD40 repeats, which fold into a β-propellor structure that provides a platform for interaction with the PTS2 motif and PTS2 co-receptor (Pan et al, 2013). While PEX5 was identified in all eukaryotic organisms, PEX7 is absent in *C. elegans*, *T. pseudonana* and *G. sulphuraria*, which may be explained by a loss of the PTS2 targeting pathway. This was shown to be the case in *C. elegans*: proteins normally containing a PTS2 have gained a PTS1 instead (Motley et al, 2000). A similar loss of the PTS2 targeting pathway has been proposed for *T. pseudonana* and the red alga *Cyanidioschyzon merolae* (Gonzalez et al, 2011). As *G. sulphuraria* is a red alga belonging to the same family as *C. merolae* (Cyanidiaceae), it is likely that the same happened in *G. sulphuraria*. Why most species utilise multiple matrix protein targeting pathways as opposed to just one is unclear. It could be that proteins of different pathways are differentially expressed depending on growth conditions, as is the case for PEX5 and its copy PEX9 in *S. cerevisiae* (Effelsberg et al, 2016, Yifrach et al, 2016). In a similar vein, it may be a matter of targeting priority, with one pathway responsible for targeting key proteins, while the other targets proteins that are less important. Another possibility is that the location of the targeting signal at either the N- or C-terminus affects protein function, making one of the targeting signals not feasible for a particular protein.

PEX5 is conserved in all eukaryotes analysed and is characterized by a disordered region at the N-terminal and several tetratricopeptide repeats (TPR) at the C-terminal (*figure 4*). While the TPR domains are responsible for its interaction with the PTS1 motif (Gatto et al, 2000), the N-terminal region interacts with a rarer PTS, PTS3 (Rymer et al, 2018) and with docking proteins PEX13 and PEX14 (Otera et al, 2002, Saidowsky et al, 2001), with the interacting regions partially overlapping (Rymer et al, 2018). As previously recognized, the structurally disordered region at the N-terminal of some PEX5 proteins shares sequence similarities with the fungi-specific PEX20 proteins (Kiel et al, 2006). These similarities between the PEX5 N-terminal and PEX20 rely on: i) a conserved motif at the N-terminal domain, ii) followed by one or more WxxxF/Y motifs and iii) a PEX7-binding domain (Schliebs & Kunau, 2006). The conserved N-terminal domain of PTS2 co-receptors contains a highly conserved cysteine residue (Schliebs & Kunau, 2006), which has been implicated in (co-)receptor recycling and cargo translocation (Hensel et al, 2011, Leon & Subramani, 2007, Okumoto et al, 2011). The WxxxY/F motifs are important for binding to PEX14 and PEX13 (Otera et al, 2002, Saidowsky et al, 2001). These WxxxF motifs are not only found in PTS2 co-receptors, but also in PEX5 of species where PEX5 does not act as PTS2 co-receptor but only as PTS1 receptor (Schliebs et al, 1999). As the name implies, the PEX7-binding domain allows the co-receptors to bind to PEX7. We checked the conservation of this domain by manually generating a hidden Markov model of the fungal PEX20, and found that this domain is detected in some but not all PEX5 orthologs that act as PTS2 co-receptors (see *figure 4*; Pex20* domains in PEX5).

**Figure 4.**
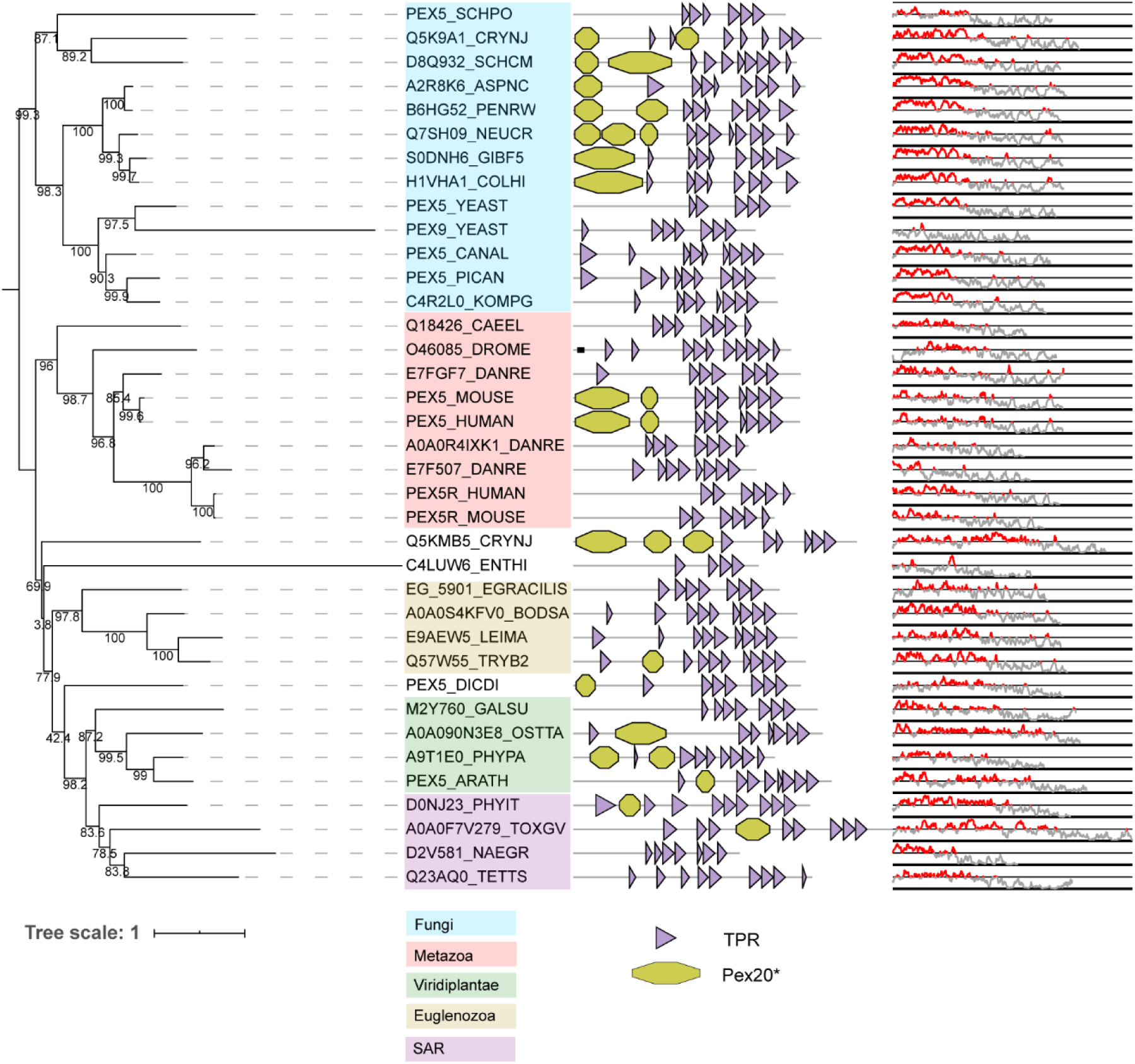
Phylogeny and protein features of PEX5 orthologs. The phylogeny is rooted at mid-point to ease the visualization and labels of the main taxonomic groups are coloured accordingly to the legend. Note that the topology does not necessarily reflect the actual evolutionary trajectory of such proteins. Protein domain architecture is defined by pfam annotations. The Pex20* is a manually generated hidden Markov model (CSM, this study). The line-dot plot indicates the regions predicted be disordered (red) and not disordered (grey).

Phylogeny shows that vertebrates and *S. cerevisiae* have duplicated their PEX5 gene independently (*figure 4*). In *S. cerevisiae,* PEX5 works as a general import receptor for all PTS1-containing peroxisomal matrix proteins, while its paralog PEX9 acts as a condition-specific receptor for a subset of PTS1 proteins (Effelsberg et al, 2016, Yifrach et al, 2016). PEX9 has lost the N-terminal disordered region that is normally present in PEX5 (see *figure 4*). Vertebrates express PEX5R, a PEX5-related protein also called TRIP8b. PEX5R is preferentially expressed in the brain and can bind PTS1- containing proteins *in vitro* (Amery et al, 2001). Nevertheless, it is unclear whether PEX5R plays any role in matrix protein targeting, although the paralogizations of PEX5 could involve different functional novelties for peroxisome protein import as in *S. cerevisiae*.

#### The docking site (PEX13 and PEX14)

Once the peroxisomal matrix protein is bound to its receptor, the receptor-cargo complex associates to the docking complex, consisting of PEX13 and PEX14 (and in fungi PEX17 or PEX33), at the peroxisomal membrane (*figure 1*).

Transmembrane helices were predicted in some, but not all, PEX13 orthologs (*figure S1A*). In addition, only in Opisthokonta organisms (fungi and metazoa) and amoebozoa, PEX13 has a predicted SH3 domain at the C-terminal (*figure S1A*), which likely controls its interaction with other proteins. PEX14 also contains a predicted transmembrane helix, but seems to be largely structurally disordered (*figure S1B)*, although it also includes several coiled-coil domains (e.g. (Lill et al, 2020)). *In vitro* protease protection experiments using human PEX13 and PEX14 confirmed that both proteins are integral membrane proteins. Human PEX14 has an N_in_-C_out_ topology, while PEX13 adopts an N_out_ -C_in_ topology, thereby exposing its SH3 domain to the peroxisomal matrix (Barros-Barbosa, Ferreira et al, 2019). The architecture of the *S. cerevisiae* PEX14-PEX17 complex was recently elucidated and revealed that PEX14 forms a 3:1 heterotetrameric complex with PEX17, forming a rod-like structure of approximately 20 nm that is exposed to the cytosol (Lill et al, 2020). This structure is mainly formed by the coiled-coil domains of PEX14 and PEX17. Besides its coiled-coil domains, PEX14 has a predicted intrinsically disordered C-terminal domain, which may be involved in recruiting import receptor PEX5 (Lill et al, 2020).

After docking, the cargo is translocated into the peroxisomal matrix. For *S. cerevisiae* PTS1 protein import it was shown that PEX5 integrates into the peroxisomal membrane to form a transient translocation pore alongside PEX14 (Meinecke et al, 2010). For PTS2 import, the pore is formed by PEX14, PEX17 and PEX18 (Montilla-Martinez et al, 2015). Little is known about the matrix protein import pores in other organisms, but the involvement of PEX14 seems to be a common denominator (Barros-Barbosa, Rodrigues et al, 2019). After formation of the translocation pore, the cargo is released into the peroxisomal matrix.

#### Receptor ubiquitination (PEX4, PEX22, PEX2, PEX10 and PEX12)

After cargo release, the PTS (co-)receptor needs to be extracted from the peroxisomal membrane, so it can be used in subsequent rounds of peroxisomal matrix protein import (Platta et al, 2014). PEX5 is mono-ubiquitinated at a conserved cysteine, leading to its extraction and recycling (Platta et al, 2014). In most eukaryotes, this ubiquitination depends on the ubiquitin-conjugating enzyme (Ubc or E2 enzyme) PEX4, associated to the peroxisomal membrane via PEX22, and on the ubiquitin ligase activities of PEX2, PEX10 and PEX12. Notably, PEX4 and PEX22 are absent in metazoa. However, mono-ubiquitination of PEX5 occurs in a comparable manner in mammalian cells through the E2D proteins UbcH5a/b/c (Grou et al, 2008). They are the closest functional counterparts to PEX4. Their actual orthologs in fungi, the Ubc enzymes, are also involved in PEX5 mono-ubiquitination (Platta et al, 2014). This reveals that in the absence of a PEX4 ortholog, functional compensation in specific organisms is possible, showing that the ubiquitination process can be shifted between subfamilies of the whole ubiquitin conjugating enzyme family. Thus, for other organisms lacking PEX4 and PEX22, it could be expected that other E2 enzymes perform this function.

The RING finger complex proteins PEX2/10/12 have ubiquitin (E3) ligase activity (Platta et al, 2014) and are broadly conserved in eukaryotes (*figure 1*). The three paralogous proteins PEX2, PEX10 and PEX12 form a heterotrimeric complex (El Magraoui et al, 2012). Characteristic for these three proteins is a highly conserved region at the N-terminus (annotated as Pex2_Pex12 pfam) and a zf-RING finger domain at the C-terminus. While the first domain can display a transmembrane helix (predicted in some of the species, suggesting membrane anchoring), the latter domain is responsible for the E3 ubiquitin ligase activity of the proteins (Platta et al, 2014) (*figure 5*). The strong conservation of both domains in most of the sequences could indicate that the cooperation of both domains is crucial for peroxisome biology. The phylogeny of these enzymes, which clearly establishes the three main subfamilies (PEX2, PEX10 and PEX12 that each contain organisms from almost all lineages), suggests that they are deep paralogs and that their functional speciation was important and early in eukaryotic evolution.

**Figure 5.**
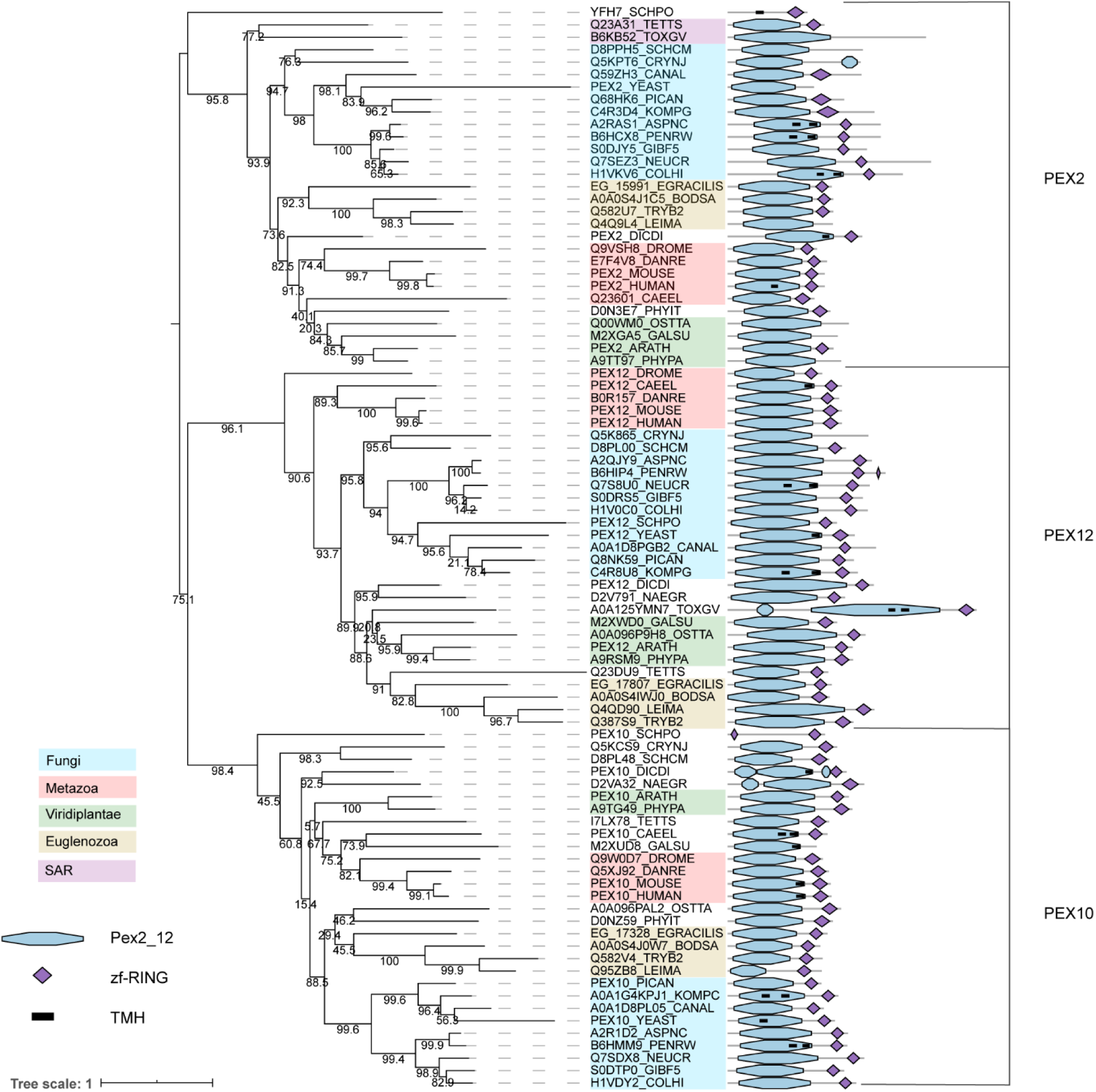
Phylogeny and protein features of PEX2/10/12 orthologs. The phylogeny is rooted at mid-point to ease the visualization and labels of the main taxonomic groups are coloured accordingly to the legend. Note that the topology does not necessarily reflect the actual evolutionary trajectory of such proteins. Protein domain architecture is defined by pfam annotations and transmembrane helix according to TMHMM software.

#### Receptor extraction (PEX1/6)

Once PEX5 is ubiquitinated, peroxisomal AAA+ ATPases PEX1 and PEX6 are responsible for PEX5 export from the peroxisomal membrane in order to recycle it back to the cytosol. PEX1 and PEX6 belong to the AAA (<u>A</u>TPase <u>a</u>ssociated with diverse cellular <u>a</u>ctivities) family (Pedrosa et al, 2018), a group of protein motors that use ATP binding and hydrolysis to mechanically unfold, disaggregate or remodel substrates (Olivares et al, 2016). Proteins of this family form ring structures with a central channel, through which they can translocate their substrates (Gates & Martin, 2020). PEX1 and PEX6 form a hetero-hexameric complex with alternating subunits in a double-ring structure (Blok et al, 2015, Gardner et al, 2015). In *S. cerevisiae*, the complex mechanically unfolds its substrates via progressive threading in an ATP-dependent manner (Gardner et al, 2015). Pedrosa et al. (2018) demonstrated using an *in vitro* setup that the PEX1/PEX6 complex directly interacts with ubiquitinated (human) PEX5, unfolding it during extraction (Pedrosa et al, 2018). The phylogeny of PEX1 and PEX6 splits both subfamilies, while their protein domain architecture shows that the architecture is more conserved in PEX1 than in PEX6 (*figure S2*). Similar to PEX2/10/12, these facts suggest that the functional speciation of PEX1 and PEX6 was also important and early in Eukaryotes.

#### The Pex11 family

Pex11 family proteins coordinate peroxisome proliferation (Koch et al, 2010). The Pex11 family is a large and complex protein family, with some members containing predicted transmembrane helices. Its phylogeny shows that it has been differentially extended in specific organisms (*figure 6*), meaning that in different lineages, independent paralogizations have occurred over time. Notwithstanding the low sequence conservation, provoking weak support in some basal nodes in the phylogeny (bootstraps lower than 80%), we can distinguish two main groups within the Pex11 protein family, which we call Pex11Y and Pex11Z here (*figure 6*). Both groups contain organisms from most taxonomic lineages, with the exception of plants, which apparently do not have Pex11Z, although they have intermediary Pex11 sequences that fall outside our Pex11Y/Z groups (along with other Pex11 protist sequences; *figure 6*). Due to the limitations of this phylogeny, it is unclear whether these forms are actually deep paralogs or whether they represent alternative evolutionary histories. However, we will follow our proposed Pex11Y/Z nomenclature to ease the functional contextualization of these paralogues.

**Figure 6.**
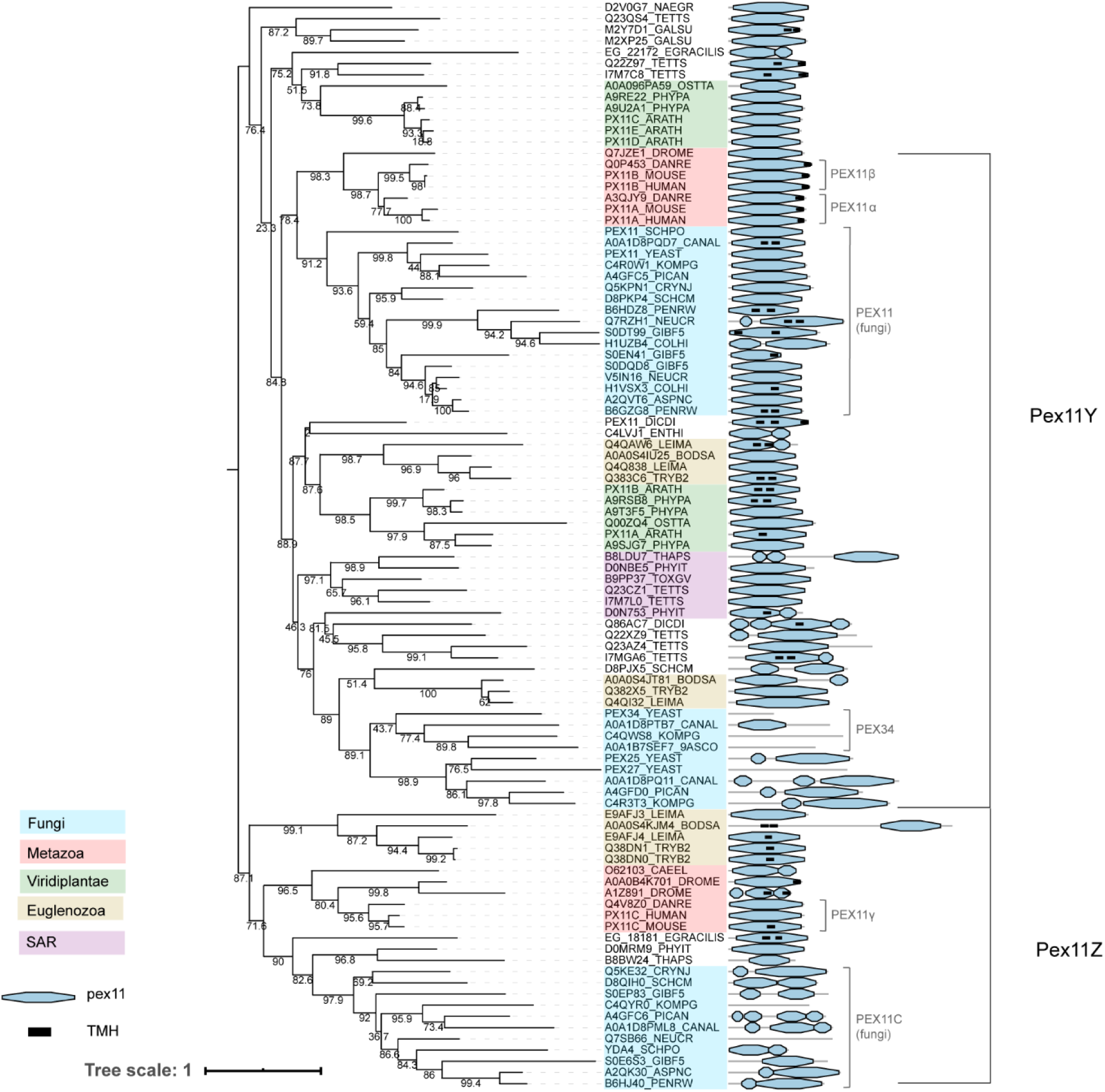
Phylogeny and protein features of PEX11 family proteins. The phylogeny is rooted at mid-point to ease the visualization and labels of the main taxonomic groups are coloured accordingly to the legend. Note that the topology does not necessarily reflect the actual evolutionary trajectory of such proteins. Protein domain architecture is defined by pfam annotations and transmembrane helix prediction (black box). The Pex11Y and Pex11Z subfamilies are named according to the most supported and basal bootstraps and their taxonomic compositions. Note that Viridiplantae organisms do not appear to have Pex11Z, although they have other paralogs outside of both defined subfamilies.

The phylogeny of PEX11 shows that these (hypothetic) deep paralogues Pex11Y/Z have subsequently undergone independent paralogizations in different lineages. For instance, Pex11Y was clearly duplicated independently in vertebrates and in several filamentous fungi. Notably, certain further paralogizations seem to have undergone extreme sequence divergence, probably providing artefactual clustering like the fungi-specific PEX25/27/34/36 subgroup within PEX11Y, which contains shortened proteins up to 144 amino acids. Fungal PEX11 and human PEX11α and PEX11β (all PEX11Y subfamily) contain a conserved amphipathic helix capable of tubulating negatively charged membranes *in vitro* (Opaliński et al, 2011, Yoshida et al, 2015). We mapped this amphipathic helix onto the multiple sequence alignment of PEX11 family proteins, observing that three positively charged residues are generally conserved in these proteins. However, the second positively charged position is not conserved in the Pex11Z subfamily (*figure S3*), suggesting possible functional difference between Pex11Y and Pex11Z. Furthermore, we observed that *S. cerevisiae* PEX34 has lost this amphipathic helix, while the C-terminal region is conserved (*figure S3*).

Several members of the Pex11 protein family have been studied. So far, the majority of studies have investigated members of the PEX11Y subfamily, which includes PEX11 from fungi and PEX11α/β from mammals. In yeasts, the absence of PEX11 results in fewer and larger peroxisomes, while cells overexpressing PEX11 have increased peroxisome numbers with smaller (Erdmann & Blobel, 1995, Joshi et al, 2012, Krikken et al, 2009). Similarly, overproduction of PEX11α or PEX11β in vertebrates induces peroxisome proliferation, while reduction of protein levels resulted in lower peroxisome numbers (Li & Gould, 2002, Schrader et al, 1998). This led to the hypothesis that these proteins play a role in peroxisome fission. Peroxisome fission takes place in three steps: organelle elongation, constriction and scission (Schrader et al, 2016). PEX11 plays a role in the first step where it functions in membrane remodelling (Schrader et al, 2016). So far, no proteins have been identified that are responsible for organelle constriction. Peroxisomal fission shares several components with the mitochondrial fission machinery, such as the dynamin related protein Dnm1 (Drp1/DLP1), Fis1 and Mff (Schrader et al, 2016). Human PEX11β recruits DRP1 to the peroxisomal membrane (Koch & Brocard, 2012, Li & Gould, 2003), and both *S. cerevisiae* PEX11 and human PEX11β have been reported to function as GTPase activating protein (GAP) for Dnm1 (DRP1) (Williams et al, 2015).

Several other functions have been attributed to proteins of the Pex11Y subfamily. *O. polymorpha* PEX11 has been implicated in peroxisome segregation during cell division (Krikken et al, 2009). *S. cerevisiae* PEX11 and PEX34 are involved in peroxisome-mitochondria contact sites (Shai et al, 2018, Ušaj et al, 2015), while *O. polymorpha* PEX11 has been implicated in peroxisome-ER contact sites (Wu et al, 2020). *S. cerevisiae* PEX11 has also been proposed to act as a pore-forming protein (Mindthoff et al, 2016) and has been implicated in medium chain fatty acid oxidation as well (van Roermund et al, 2000). As only a subset of proteins from the PEX11Y subfamily have been investigated, perhaps other functions will still be discovered.

Much less is known about proteins of the PEX11Z subfamily, which includes PEX11γ from metazoa, fungal PEX11C and GIM5A/B from *T. brucei*. However, they also play a role in peroxisome proliferation (see e.g. (Koch & Brocard, 2012, Opaliński et al, 2012)). PEX11γ has been suggested to coordinate peroxisomal growth and division via heterodimerisation with other mammalian PEX11 paralogs and interaction with Mff and Fis1 (Schrader et al, 2016). *O. polymorpha* PEX11C is downregulated upon shifting from peroxisome repressing (glucose) to peroxisome inducing (methanol) growth conditions (van Zutphen et al, 2010) suggesting that PEX11C is not required for peroxisome proliferation. In *Penicillium rubens*, deletion of PEX11C has no significant effect on peroxisome number or size, while overexpression strongly stimulates peroxisome proliferation (Opaliński et al, 2012). In *T. brucei*, the absence of both GIM5A and GIM5B is fatal, due to cellular fragility (Voncken et al, 2003). In *S. cerevisiae* proteins of the PEX11Z subfamily are absent.

The remaining proteins, which do not have a clear evolutionary relationship with each other and fall outside the Pex11Y/Z subfamilies, we call the Pex11-like proteins. The most studied proteins from this group are *A. thaliana* PEX11C/D/E. These proteins cooperate with FIS1b and DRP3A in peroxisome growth and division during the G_2_ phase just prior to mitosis (Lingard et al, 2008). Interestingly, in cells where PEX11C, PEX11D and PEX11E were silenced simultaneously, peroxisomes were enlarged, but not elongated, suggesting that these proteins act in peroxisome growth, but not tubulation (Lingard et al, 2008).

### 3. PEX proteins specific for fungi

Several PEX proteins are specific to fungi. The high number of known fungal PEX proteins is probably due to the extensive screens for yeast peroxisome-deficient mutants that have been performed in the past (Erdmann et al, 1997). Additionally, current peroxisome biogenesis research is still taking advantage of a wealth of genetic and biochemical toolboxes to analyse the molecular biology of these organelles in yeast.

#### The PEX7 co-receptors (PEX18, PEX20, PEX21)

In plants, animals and protists like TRYPB2, *D. discoideum* and *L. major*, (a longer splicing variant of) PEX5 acts as PEX7 co-receptor for PTS2 protein import (Schliebs & Kunau, 2006). In contrast, in many fungi the PEX7 co-receptor is a separate PEX protein, namely PEX18, PEX20 or PEX21 (see for more detailed reviews e.g. (Kunze, 2020, Schliebs & Kunau, 2006)). Duplication of the ancestral PEX20 in *S. cerevisiae* (see *figure S4*), resulted in the partially redundant paralogs PEX18 and PEX21 that perform the same function (Purdue et al, 1998). Therefore, these proteins can be considered as a single PEX20 group. As previously described, some sequence features relate PEX20 with the N-terminus of PEX5 proteins: a conserved cysteine, WxxxF motifs and PEX7 binding domain (Schliebs & Kunau, 2006). Due to the fact that that PEX5 is present in most eukaryotes and Pex20 domains can be found at the N-terminus of many such proteins, it is most likely that PEX20 is the result of a protein domain separation specific to fungi, rather than the previously proposed protein fusion of PEX5 and PEX20 (Kiel et al, 2006).

#### PEX17 and PEX33

In all species, PEX13 and PEX14 are components of the receptor docking site. An additional component of the docking site in yeasts is PEX17, while in filamentous fungi PEX33 is part of the docking complex. PEX17 is characterized by a single transmembrane helix at the N-terminal. As described above, *S. cerevisiae* PEX14 and PEX17 together form a rod-like structure at the peroxisomal membrane (Lill et al, 2020). PEX33 is a paralog of PEX14, whereas PEX17 is a protein partially aligning to the C-terminal of PEX14 and PEX33, suggesting PEX17 is a PEX14-like protein. The exact functions of PEX17 and PEX33 are still unclear, but PEX17 in *S. cerevisiae* is a main component of the PTS2 import pore (Montilla-Martinez et al, 2015) and seems to increase the efficiency of binding of import receptors PEX5 and PEX7 to the docking complex (Lill et al, 2020).

#### PEX8

In fungi, intraperoxisomal protein PEX8 bridges the docking and RING finger complexes (PEX2/10/12) (Agne et al, 2003). Little else is known about PEX8, but it has been implicated in cargo release from the PTS1 receptor PEX5 (Ma et al, 2013, Wang et al, 2003).

#### Pex23 family proteins

PEX23, PEX24, PEX29, PEX32 (for *O. polymorpha* for example) and PEX28, PEX29, PEX30 PEX31, PEX32 (for *S. cerevisiae*) are homologous proteins containing a highly conserved domain called the Pex24p domain (pfam). This domain contains a Dysferlin (DysF) motif at the C-terminal region, the function of which is still unclear (Wu et al, 2020). At the N-terminal, these proteins have several transmembrane domains suggesting that these proteins are anchored to membranes. A group of proteins related to this Pex23 protein family are the Pex23-like proteins (Kiel et al, 2006), including SPO73, a protein involved in sporulation. Pex23-like proteins do not usually present the region containing the predicted transmembrane helices. The phylogeny of all these proteins can be divided into three main groups that here we call PEX23 subfamily, PEX24 subfamily and Pex23-like proteins (*figure 7A*). The sequences from the PEX23 and PEX24 subfamilies appear to differ in protein extensions at their C- and N-termini respectively, with predicted structural protein disordered regions. Due to the fact that the main PEX23 and PEX24 subfamilies contain most of the fungi analysed, it is likely that both subfamilies originated from an ancestral duplication in fungi. Later, these PEX23 and PEX24 paralogs duplicated in yeasts leading to amongst others PEX28/PEX29 and PEX30/31/32 in the ancestor of *S. cerevisiae.* In filamentous fungi on the other hand, no duplication occurred, and these fungi express only one protein of each group. Thus, these proteins have diversified differentially in Fungi.

**Figure 7.**
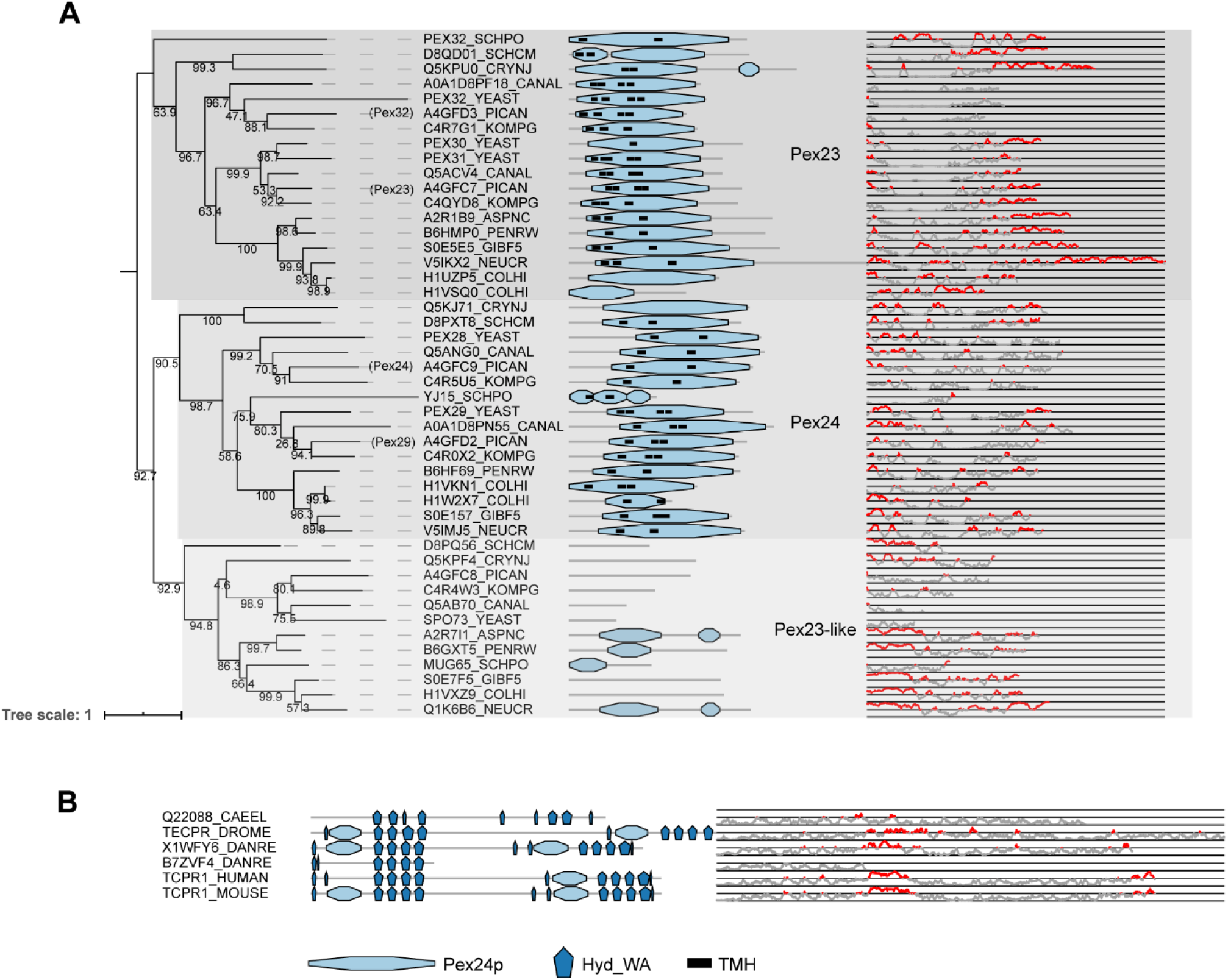
Phylogeny and protein features of Pex23 protein family and TECPR1 proteins. A) Phylogeny and protein features of the fungal Pex23 protein family. The phylogeny is rooted at mid-point to ease the visualization. The main phylogenetic groups are named and highlighted according the protein names of *O. polymorpha*, indicated between brackets. Protein domain architecture is defined by pfam annotations and transmembrane helix according to TMHMM software. The Pex24p pfam domain contains the DysF motifs. The line-dot plot, indicates the region predicted be disordered (red) and not disordered (grey). B) Protein features of TECPR1 protein family from metazoa.

#### Unlike other peroxins, proteins of the Pex23 family localise to the ER instead of peroxisomes

Although initially reported at the peroxisome (Brown et al, 2000, Tam & Rachubinski, 2002, Vizeacoumar et al, 2003, Vizeacoumar et al, 2004), later studies either reported dual localization to peroxisomes and ER (David et al, 2013, Yan et al, 2008) or exclusive localization at ER subdomains (Joshi et al, 2016, Mast et al, 2016, Wu et al, 2020). A recent study characterizing *O. polymorpha* Pex23 family members reported the involvement of PEX24 and PEX32 in peroxisome-ER contact sites (Wu et al, 2020). This could explain the previous contradictory reports on their localization, as they can be expected to be present in spots where peroxisomes and ER interact. Furthermore, *S. cerevisiae* PEX30 and PEX31 are ER membrane shaping proteins (Joshi et al, 2016). *S. cerevisiae* PEX30 plays a role in regulating budding of pre-peroxisomal vesicles and lipid droplets from specific ER subdomains (Joshi et al, 2016, Joshi et al, 2018). It has been proposed to facilitate this by collaborating with seipin to organize ER subdomains to alter the membrane lipid composition (Wang et al, 2018). In humans, no orthologs of PEX30 have been identified, but MCTP2 has been suggested to act as a functional analog (Joshi et al, 2018).

#### PEX23 homologs were found in metazoa, but these proteins cannot be considered orthologs of PEX23

These proteins were previously published as metazoan PEX23 orthologs (e.g. (Di Cara et al, 2017, Jeynov et al, 2006, Mast et al, 2011)) and are also annotated as such in some databases (e.g. protein Q9VWB0|TECPR_DROME annotated as PEX23 in Uniprot and FlyBase). However, their domain architecture (see *figure 7B*) is clearly different from previously established Pex23 family proteins, and they actually belong to the TECPR1 family of proteins. TECPR1 proteins are localized to lysosomes and play a role in autophagy (Chen & Zhong, 2012). While TECPR1 proteins do contain a DysF domain, like the proteins from the PEX23 family, they also contain several tectonin repeats (TECPR) and a PH domain, in addition to a beta-propellor structure (Ogawa et al, 2011). It is therefore unlikely that they perform a function similar to PEX23 family proteins in fungi and they cannot be considered PEX23 orthologs.

#### Little is known about both PEX35 and PEX37, but both seem to play a role in regulating peroxisome proliferation

PEX35 is unique to *S. cerevisiae* and closely related species in the Saccharomycetaceae family, while PEX37 is found in most other yeast species and filamentous fungi. PEX35 has no known functional domains or similarity to other known PEX proteins. Only one study investigating PEX35 has been published to date, showing that PEX35 is a PMP that interacts with vesicle budding inducer Arf1 and localizes at the proximity of proteins from the Pex11 family (Yofe et al, 2017). The authors speculate that PEX35 may regulate peroxisome fission alongside proteins of the Pex11 family. *O. polymorpha* PEX37 is a peroxisomal transmembrane protein that affects peroxisome segregation and proliferation under peroxisome-repressing conditions, but not on peroxisome-inducing conditions. So far, only one study has investigated this protein (Singh et al, 2020). PEX37 belongs to the same protein family as human PXMP2, *N. crassa* Woronin body protein Wsc and *S. cerevisiae* mitochondrial inner membrane protein Sym1 and its human homolog MPV17, many of which are thought to act as channels. Human PXPM2 is able to partially rescue the phenotype present in the absence of *O. polymorpha* PEX37, suggesting that these proteins have similar functions (Singh et al, 2020).

### 4. Moderately conserved PEX proteins

In many species, the PEX1/PEX6 complex is recruited to the peroxisomal membrane via an anchoring protein. These membrane anchors are much less conserved than PEX1 and PEX6 themselves, with different homologous, but not orthologous, proteins acting as anchoring protein in different species. In vertebrates and most fungi, the anchoring protein is PEX26, while in *S. cerevisiae* and closely related species in the Saccharomycetaceae family it is PEX15 (Kiel et al, 2006) and in plants it is APEM9 (Cross et al, 2016). Despite sharing only weak sequence identity, PEX15, PEX26 and APEM9 do have several features in common. All three proteins are tail-anchored proteins (Cross et al, 2016, Halbach et al, 2006) and tether the PEX1/PEX6 complex to the peroxisomal membrane via PEX6 (Birschmann et al, 2003, Goto et al, 2011, Matsumoto et al, 2003).

## Discussion

We used a comparative genomics approach to provide an up-to-date overview of all PEX protein families known to date, in a range of representative organisms from all eukaryotic lineages. Our computational survey identified a **core set of PEX proteins** that is broadly conserved across all eukaryotic lineages (PEX1/2/3/5/6/7/10/11/12/13/14/16/19). This means that ancestral versions of these PEX proteins were already present in the last eukaryotic common ancestor (LECA) and that this set of proteins defines the minimum set of PEX proteins that is required to make a peroxisome. Besides a broadly conserved core set of PEX proteins, we found that a large number of **PEX proteins is specific to the kingdom of fungi**. Although there is increasing consensus that homology detection failure is frequent (Weisman et al, 2020), our inner controls (see methods) still suggest that these fungi-specific proteins are absent in other lineages. This exposes not only a bias in peroxisome research towards fungi, but also reveals that peroxisomes are dynamic organelles, their composition evolving under different evolutionary pressures. The loss of specific PEX proteins in some eukaryotes, such as the loss of proteins associated with the PTS2 targeting pathway in *C. elegans* and the loss of PEX16 in *S. cerevisiae* and other yeasts further supports this notion.

Intriguingly, PEX proteins in human pathogens like *T. gondii*, *T. brucei* and *L. major* were often difficult to detect. Moreover, these PEX proteins frequently had additional domains, which could indicate that they may have obtained additional functions. The low homology between PEX proteins of human and human pathogens may be advantageous for the identification of specific drug targets.

In some species lacking peroxisomes such as *E. histolytica*, we still identified some PEX proteins such as PEX5, PEX16 and PEX19 (see *figure 2, 3* and *4*). This could mean that these organisms have lost this organelle relatively recently and thus have not entirely lost all PEX proteins yet, but it could also suggest that the remaining PEX proteins retain non-peroxisomal functions. This is not completely unthinkable, as some PEX proteins have already been suggested to be involved in non-peroxisomal pathways. For instance, human PEX3 and PEX19 have been implicated in targeting of lipid droplet protein UBXD8 (Schrul & Kopito, 2016). On the other hand, the example of *E. histolytica* illustrates drastic evolutionary changes in the peroxisomal biology, a fact already observed in other amoeba species like *Mastigamoeba balamuthi* (Le et al, 2020). Understanding the reason behind these evolutionary adaptations will improve our understanding about the peroxisome biology.

The vast majority of the core PEX proteins (PEX1/2/5/6/7/10/12/13 and 14) are involved in matrix protein import, while only a few (PEX3, PEX16 and PEX19) play a role in PMP sorting. In addition to these core PEX proteins all eukaryotes contain multiple proteins of the Pex11 family, which are involved in several peroxisome-related processes. It is unclear why so few proteins have been identified that plays a role in PMP sorting. Proteins of the common ER protein sorting machineries, such as the Sec and GET translocons, have been reported to function in the indirect pathway of PMP sorting. The absence of these proteins is lethal in yeast, explaining that such mutants have not been obtained in screens for yeast peroxisome deficient mutants. For the direct pathway of PMP sorting it is unlikely that the entire sorting/insertion machinery consists of only three, or even two for yeast (PEX3/19), proteins.

Most of the currently known PEX genes have been identified in the nineties of the previous century by very successful genetic approaches to identify peroxisome deficient (*pex*) yeast mutants. Yeast *pex* mutants are viable and have distinct growth phenotypes (e.g., deficiency to grow on oleic acid or methanol), which greatly facilitated the isolation of these mutants and cloning of the corresponding genes by functional complementation. Most likely this caused the bias towards fungal PEX genes. In addition to *S. cerevisiae*, which is the main yeast model in cell biology, a few other yeast species were used to identify PEX proteins (*Komagataella phaffiia* [formerly *Pichia pastoris*], *Ogataea polymorpha* [formerly *Hansenula polymorpha*] and *Yarrowia lipolytica*). Notably, several conserved PEX proteins that are present in the latter three yeast species are absent in *S. cerevisiae* (for instance PEX20, PEX26, PEX37 and proteins of the PEX11Z subfamily), while orthologs of the *S. cerevisiae* PEX proteins PEX9, PEX15, PEX35 are absent in all other species that we analysed. This stresses the importance of using several yeast models besides *S. cerevisiae* in cell biology research.

Fusion of human cell lines, derived from patients suffering from peroxisome biogenesis disorders, resulted in the classification of these patients in 12 genotypes/complementation groups (Fujiki, 2016). Using known yeast PEX genes, human orthologues were identified by homology searches on the human expressed sequence tag database. By functional complementation of the cell lines with these putative human PEX genes, 12 of the currently known human PEX genes were identified. Because mislocalisation of the PTS1 protein catalase was used as criterion for peroxisome deficiency, the human PTS2 receptor PEX7 was not identified by this approach (Fujiki, 2016). Together with the results of functional complementation of mutant Chinese hamster ovary (CHO) cell lines, at present 16 mammalian PEX proteins are known (compared with 29 in *S. cerevisiae*).

It is unlikely that all human/mammalian PEX proteins have been identified. Mutations in human/mammalian PEX genes could cause lethal phenotypes, explaining why they have not been isolated in mutant screens. Also, there may be functional redundancy among human PEX genes, which prevents their identification by mutant complementation approaches. Conversely, mutations in yet unknown mammalian PEX genes could cause relatively weak phenotypes and hence were overlooked. Indeed, the approaches used so far resulted in the identification of PEX11β, but not of PEX11α and PEX11γ. Alternative approaches, like the identification of novel peroxisomal proteins using proteomics of isolated mammalian peroxisomes may result in the characterization of novel mammalian PEX proteins.

PEX proteins (peroxins) were originally defined as proteins “involved in peroxisome biogenesis (inclusive of peroxisomal matrix protein import, membrane biogenesis, peroxisome proliferation, and peroxisome inheritance)” (Distel et al, 1996). However, proteins fitting this definition are not always named as such. For instance, *T. brucei* GIM5A is a member of the Pex11 protein family, but is not named ‘PEX’. Also, two proteins involved in peroxisome inheritance, Inp1 and Inp2, are not called PEX. Therefore “inheritance” could be omitted from the original definition of PEX proteins, or these proteins could be renamed. Some proteins that fulfil the PEX protein definition are also involved in other processes and obviously not called PEX. This is for instance the case for the organelle fission proteins FIS1 and DRP1, and ER proteins that play a role in the indirect sorting pathways of PMPs.

### Current PEX protein nomenclature has several issues and inconsistencies that can easily lead to confusion

As PEX proteins are numbered chronologically, there is no intuitive link between their names and their function and/or conservation. Additionally, there are several naming inconsistencies relating to PEX protein families. For instance, in higher eukaryotes Pex11 protein family members are named PEX11‘X’ (e.g., PEX11α/β/γ, PEX11A/B/C). The nomenclature of yeast proteins does not allow the addition of the extra symbol ‘X’. These genes invariably consist of a three-letter code (PEX) followed by a number, explaining why PEX11 orthologs in yeast have been designated PEX25, PEX27 and PEX34, not PEX11X.

Since most PEX proteins were initially identified in yeast species and numbered in the order in which they were described, proteins belonging to the same protein family have received different names. For instance, the two AAA ATPases are called PEX1 and PEX6, while the three RING proteins are called PEX2, PEX10 and PEX12. Lastly, there are proteins carrying the same name that are not actually orthologs (e.g., PEX23 in metazoa). In summary, current PEX protein nomenclature can easily lead to confusion as it is often far from intuitive, sometimes inconsistent and occasionally wrong. This not only leads to confusion within the peroxisome field, but the large number of PEX proteins numbering up to 37 can be quite intimidating for researchers from other fields.

We therefore suggest that it may be prudent to come up with a **new naming system**. Although it is beyond the scope of the current paper, similar new naming systems are not unprecedented. Indeed, the name PEX protein itself was devised to unify nomenclature regarding proteins involved in peroxisome biogenesis (Distel et al, 1996), thereby re-naming the 13 proteins known at the time to be involved in peroxisome biogenesis. More recently, proteins involved in mitochondrial contact site and cristae organizing system (MICOS) (Pfanner et al, 2014), autophagy-related proteins(Klionsky et al, 2003) and ribosomal proteins (Ban et al, 2014) have been re-named. In addition, we recommend setting up guidelines for naming newly discovered ‘PEX proteins’, taking into account phylogeny to extend to ortho- and in-paralogues. Moreover, we propose amending the definition of ‘PEX proteins’ as posed in 1996 (Distel et al, 1996). Proteins involved in peroxisome inheritance such as Inp1 and Inp2 have so far been named differently and should be removed from the definition.

Adopting an entirely new naming system may be very difficult. However, it would already be very helpful to only re-name the most confusing and inconsistent parts. The two largest protein families, the Pex11 family and the Pex23 family, together make up about one-third of all PEX numbers and are arguably the most confusingly named.

## Acknowledgements

RLMJ is supported by a grant from the Netherlands Organization of Scientific Research (NWO), section Earth and Life Sciences (ALWOP161). DPD was supported by the Spanish Ministry of Economy and Competitiveness (Grant nos. BFU2016-78326-P). CSM is supported by the Joined Gordon and Betty Moore Foundation with the Simons foundation (Grant Agreement #9733).

## Author contributions

RLMJ, CSM, MVDN, DPD and IvdK conceived the project; RLMJ, CSM, MVDN, DPD analysed the data and prepared the figures; RLMJ and IJvdK wrote the original draft. All contributed to reviewing and editing the manuscript.

## Conflict of interest

The authors declare no conflict of interest.

## Methods

### Ortholog identification of PEX proteins

For the ortholog detection of PEX proteins, we systematically used two approaches: reciprocal searches of single protein sequences and reciprocal searches based on protein profiles (Hidden Markov models). We selected a set of eukaryotic proteomes from UniProt (Anonymous, 2017) (see *table 1*) and for both approaches, performed the reciprocal searches starting from the sequences of different organisms (see table) and made a consensus for the assignment of orthologs between the searches.

The first approach was based on phmmer searches (HMMER package (Potter et al, 2018)). As peroxisomal proteins can be multidomain proteins, when the first reciprocal hit failed, we also checked the best domain e-value hit from the target proteome. In this way, we also retrieve potential orthologs taking into account alternative domain architecture. The second approach was based on reciprocal jackhmmers followed by hmmsearches (HMMER package (Potter et al, 2018)). This method is applied in order to detect divergent orthologs undetectable by the previous approach, although it can be problematic for proteins containing common domains like PEX1/6, PEX4 (containing functional domains like WD40, ATPase, zinc-finger and ubiquitin ligases; see table). Due to the diverse nature of the PEX proteins, different e-value thresholds and iterations were applied. For example, searches involving transmembrane proteins and tandem protein repeats (TPR) were conducted with 2 iterations and a relaxed e-value, 1e-2. Alternatively, for the other common domains, we applied 2 iterations and constrained e-value, 1e-20. The reciprocal detection for these common domains were often/frequently unsatisfying showing the limitation of this method for abundant and common domains.

Once the ortholog assignment of both methods combined, for each set of orthologs we manually filtered-out possible false positive by performing a multiple sequence alignment using Mafft (einsi-mode (Katoh & Standley, 2013)) followed by visual inspection. We additionally searched for missing orthologs. We built HMM profiles through Hmmbuild using the MSA generated previously and made searches into the suspect proteome through Hmmsearch (both from the HMMER package (Potter et al, 2018)). It is important to note that if no orthologs were identified for a particular PEX protein in a specific organism, this does not necessarily mean that no ortholog exists. Possible causes of not identifying orthologs are incomplete genome information and sequence divergence of the ‘true’ ortholog. For example, the *T. pseudonana* proteome seems to be incomplete in the Uniprot database: a previous study identified a *T. pseudonana* Pex12 ortholog (Mix et al, 2018) that matches our criteria for orthology, but is absent from Uniprot.

Ortholog sequences included in the final dataset were aligned with Mafft, and trimmed the gap position with Trimal using different thresholds. Phylogenetic trees were constructed using IQ-TREE (Nguyen et al, 2015) obtaining branch supports with ultrafast bootstrap (Hoang et al, 2018) and applying the automatic model selection calculated by ModelFinder (Kalyaanamoorthy et al, 2017). Trees were visualized and annotated using iTOL (Letunic & Bork, 2019). Functional domain annotation was carried out using the Pfam database (El-Gebali et al, 2019), transmembrane domains using the TMHMM server (http://www.cbs.dtu.dk/services/TMHMM/) and structural disorder with IUPred2 (Mészáros et al, 2018).

## Supplementary information

**Figure S1:**
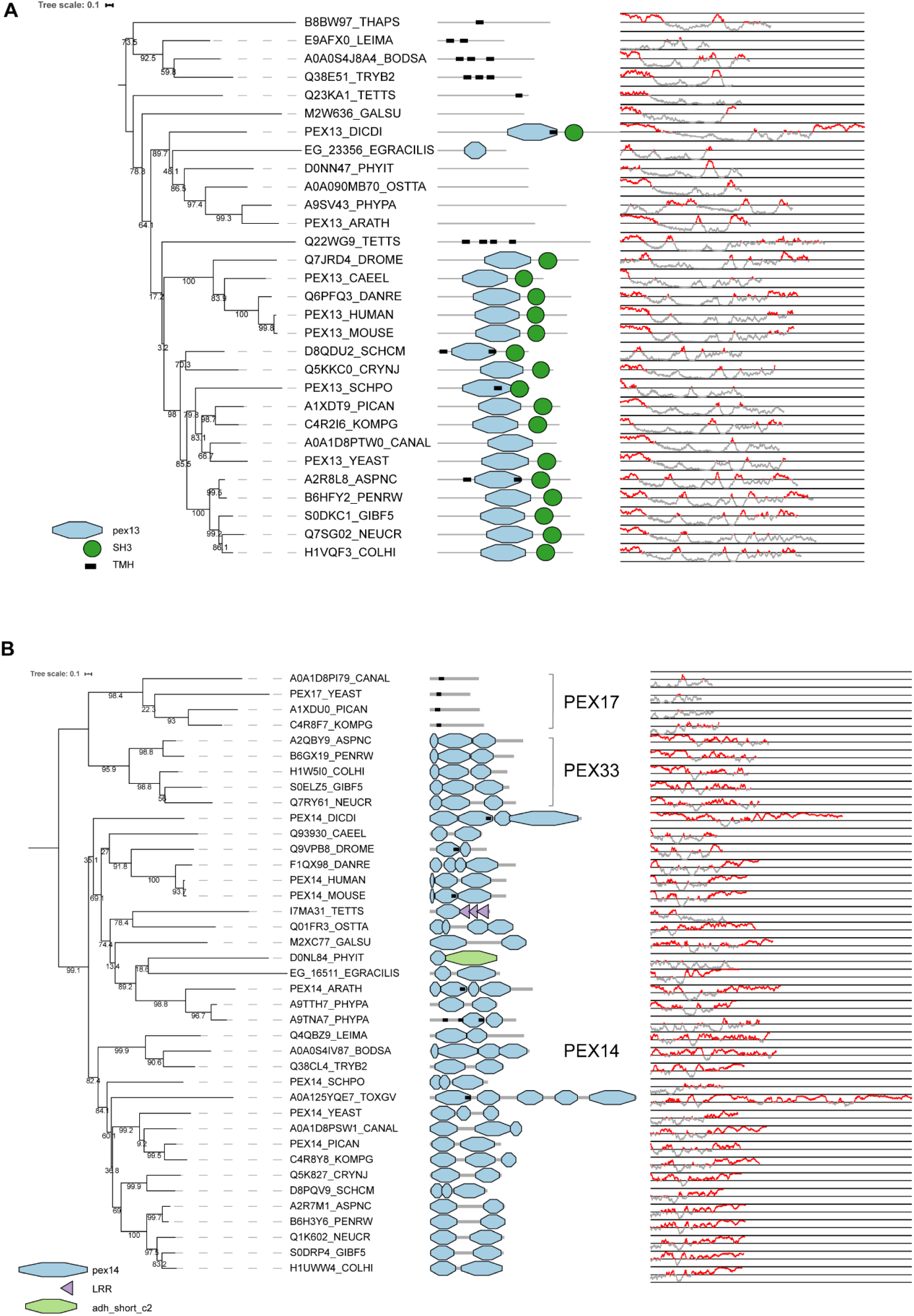
Phylogeny and protein features of A) PEX13 and B) PEX14/17/33 orthologs. The phylogeny is rooted at mid-point to ease the visualization. Note that the topology does not necessarily reflect the actual evolutionary trajectory of such proteins. Protein domain architecture is defined by pfam annotations and transmembrane helix according to TMHMM software. The line-dot plot, indicates the regions predicted to be disordered (red) and not disordered (grey).

**Figure S2:**
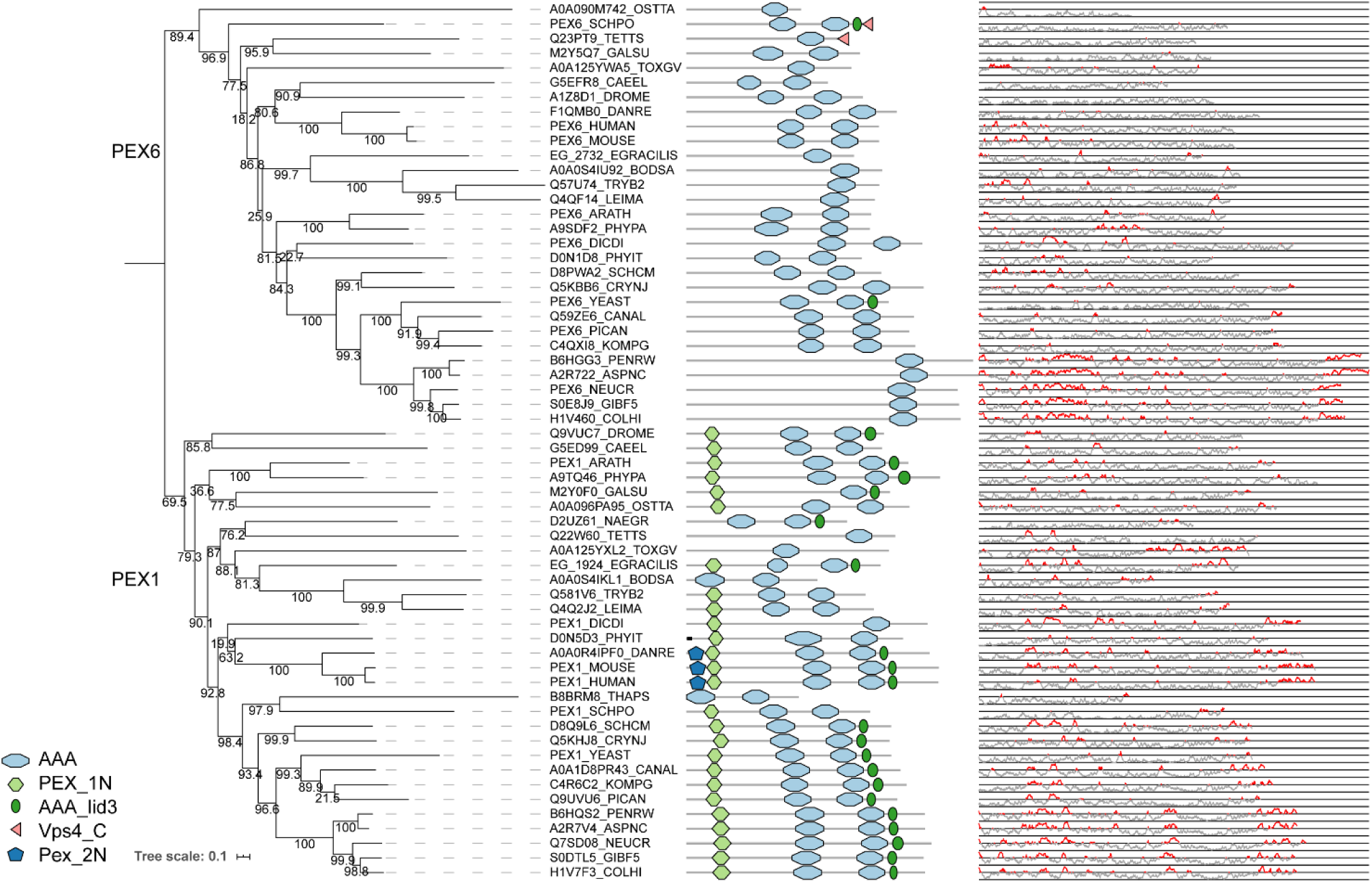
Phylogeny and protein features of PEX1/6 orthologs. The phylogeny is rooted at mid-point to ease the visualization. Note that the topology does not necessarily reflect the actual evolutionary trajectory of such proteins. Protein domain architecture is defined by pfam annotations and transmembrane helix according to TMHMM software. The line-dot plot, indicates the regions predicted to be disordered (red) and not disordered (grey).

**Figure S3:**
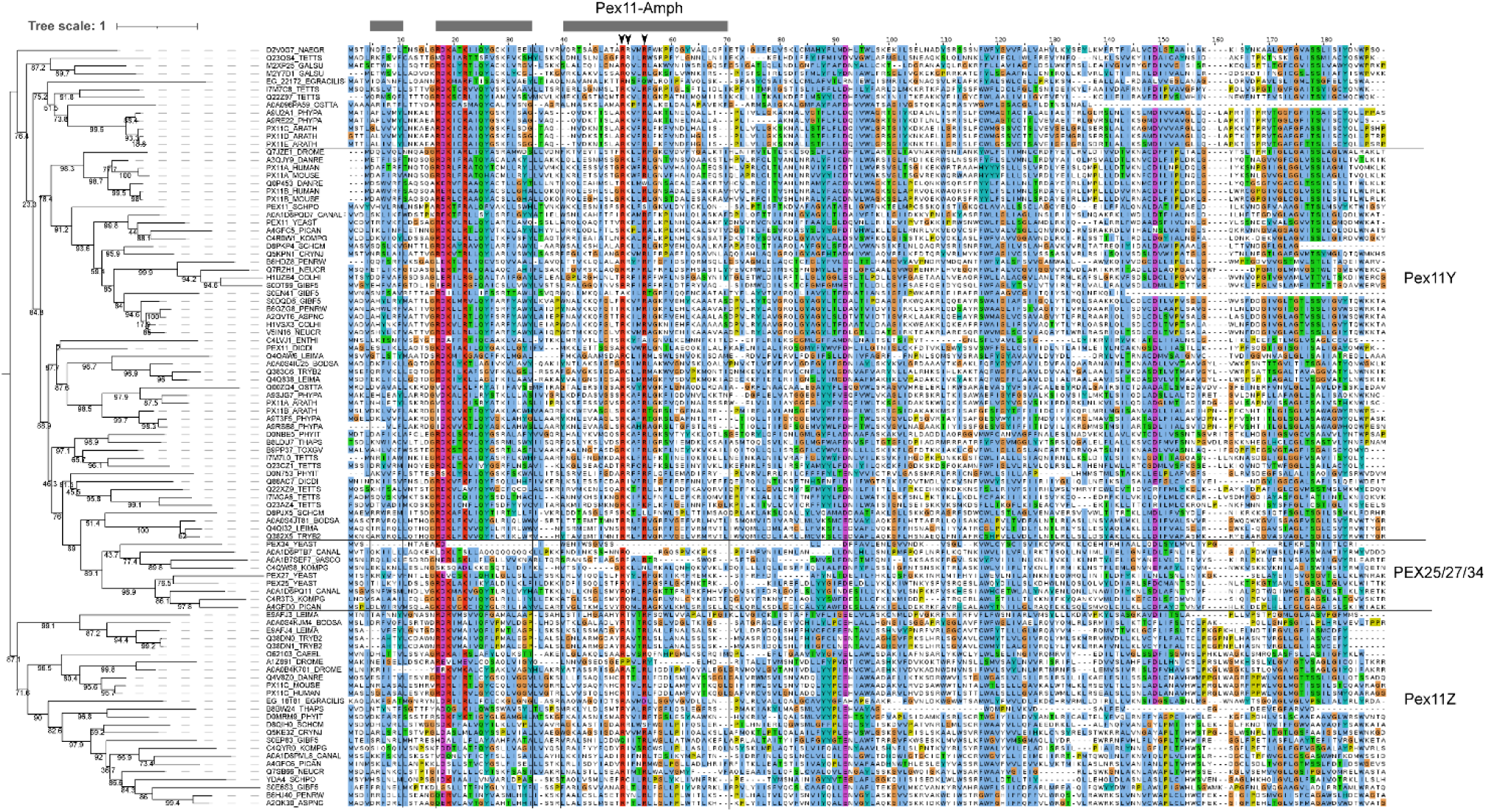
Multiple sequence alignment of Pex11 family proteins. Grey bars above sequences denote predicted α-helices and the N-terminal amphipathic helix (Pex11-Amph). Residues are coloured based on physico-chemical properties according to ClustalW. The phylogeny is rooted at mid-point to ease the visualization. Note that the topology does not necessarily reflect the actual evolutionary trajectory of such proteins.

**Figure S4:**
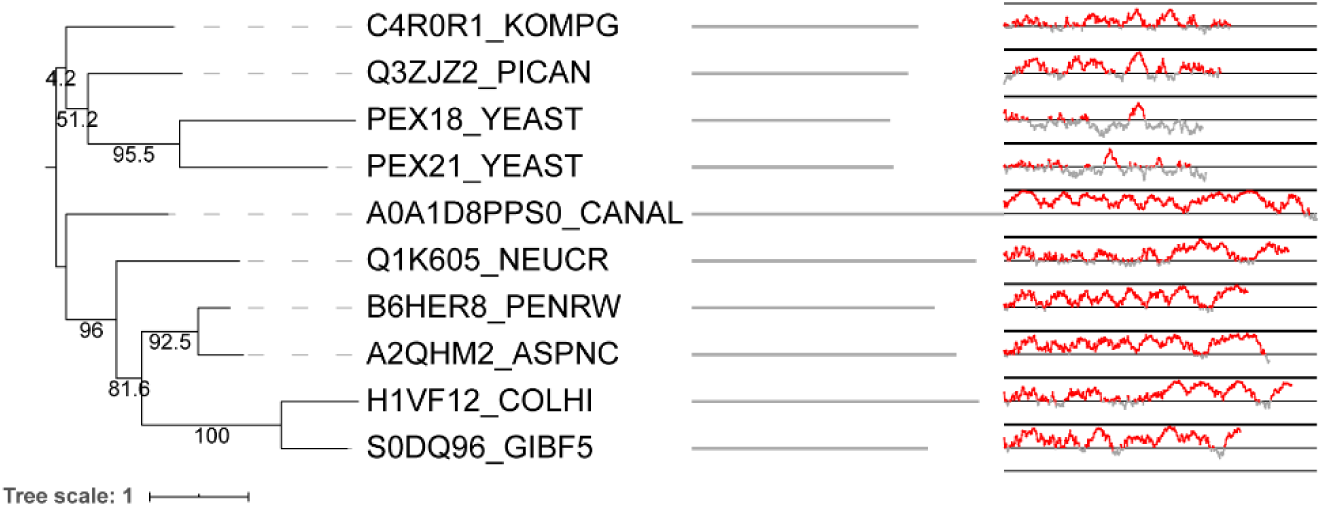
Phylogeny and protein features of PEX118/20/21 orthologs. The phylogeny is rooted at mid-point to ease the visualization and labels of the main taxonomic groups are coloured accordingly to the legend. Note that the topology does not necessarily reflect the actual evolutionary trajectory of such proteins. Protein domain architecture is defined by pfam annotations and transmembrane helix according to TMHMM software. The line-dot plot, indicates the regions predicted to be disordered (red) and not disordered (grey).

**Table S1:**
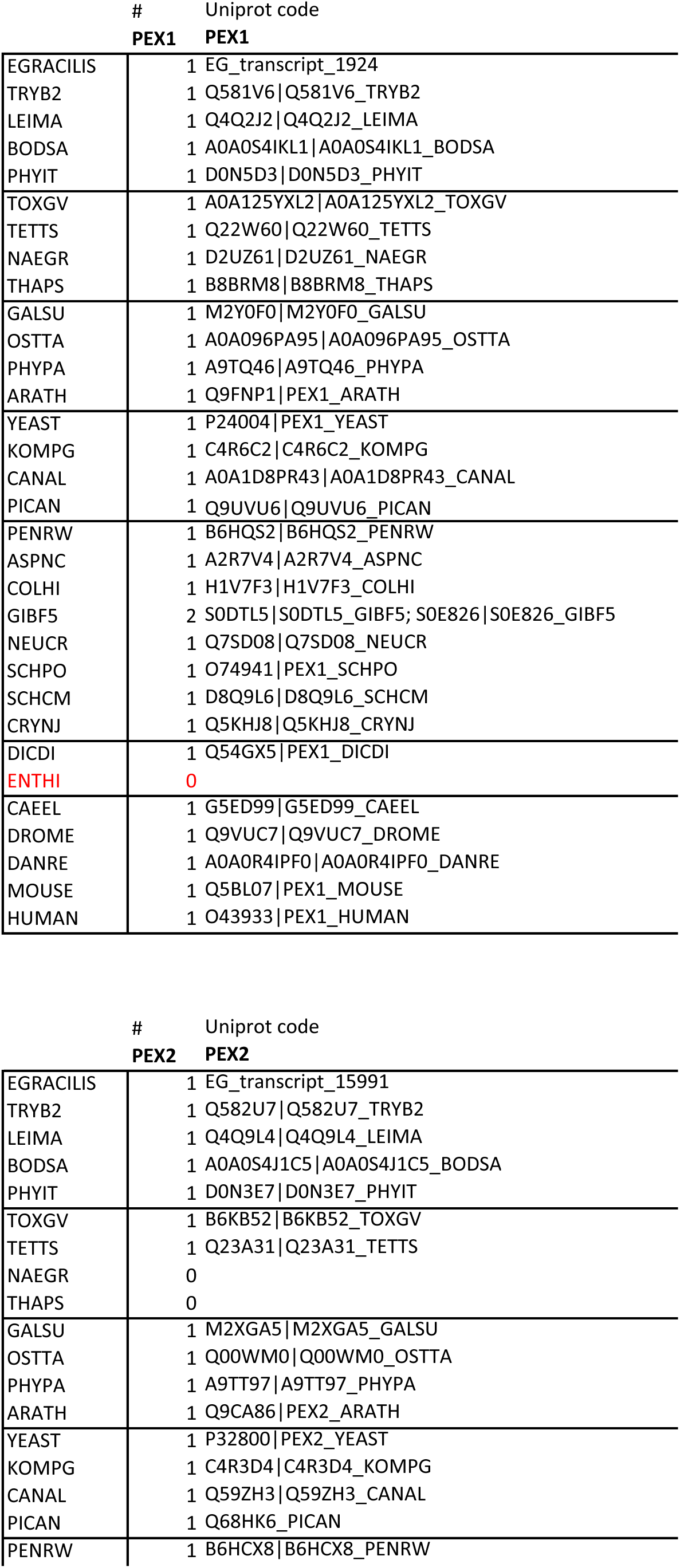
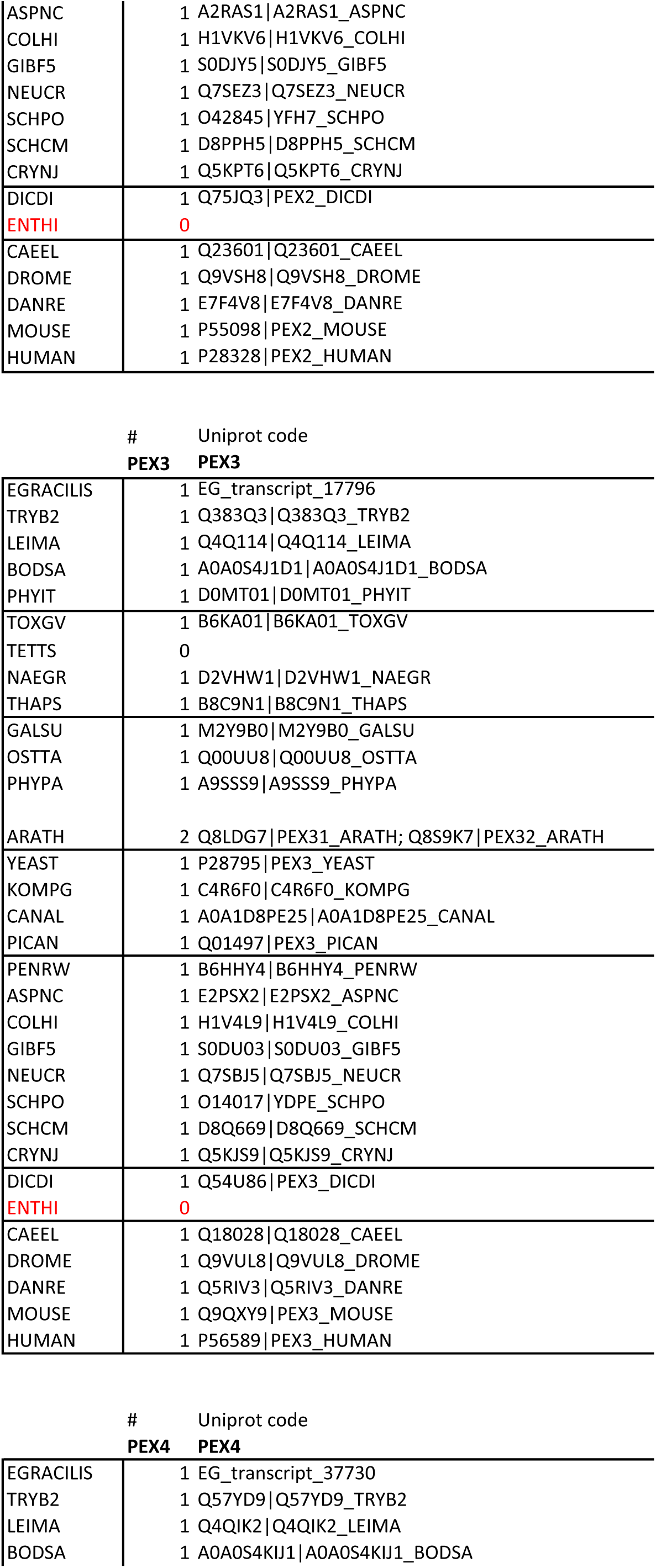
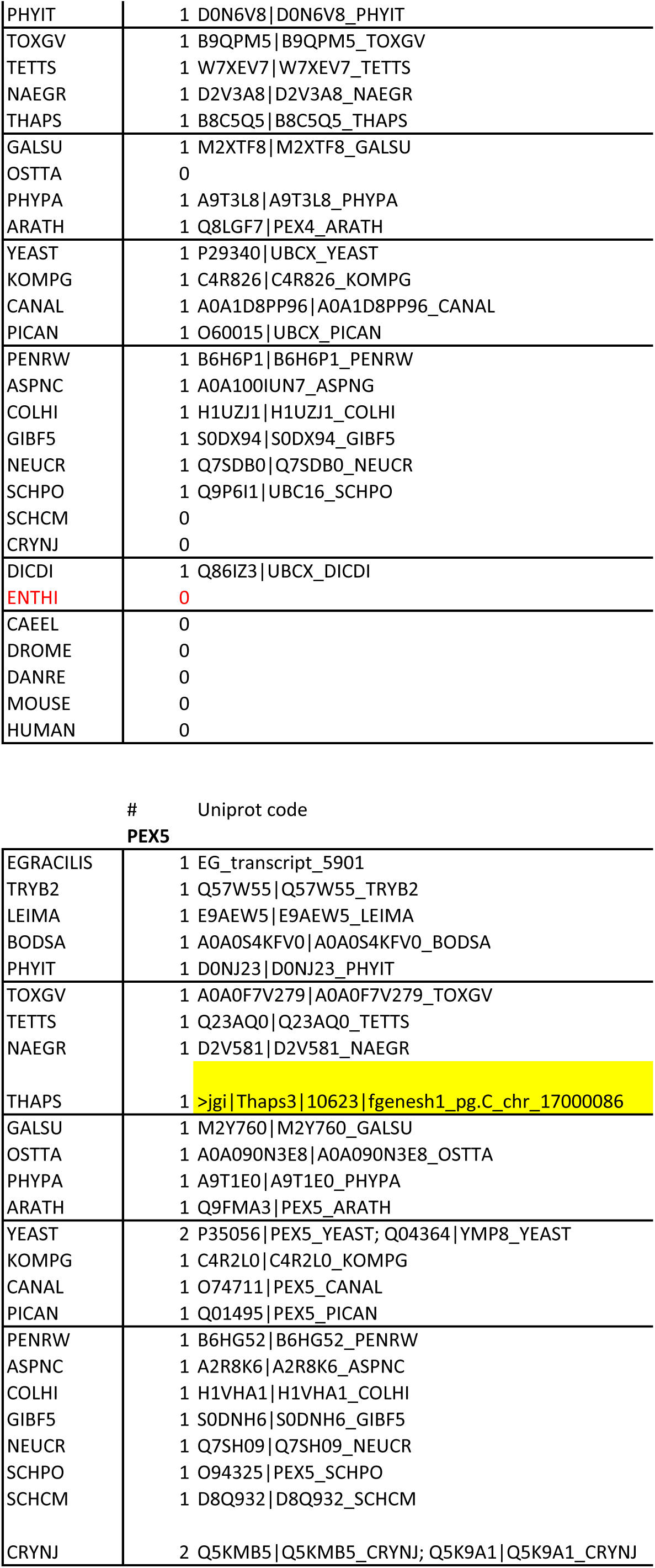
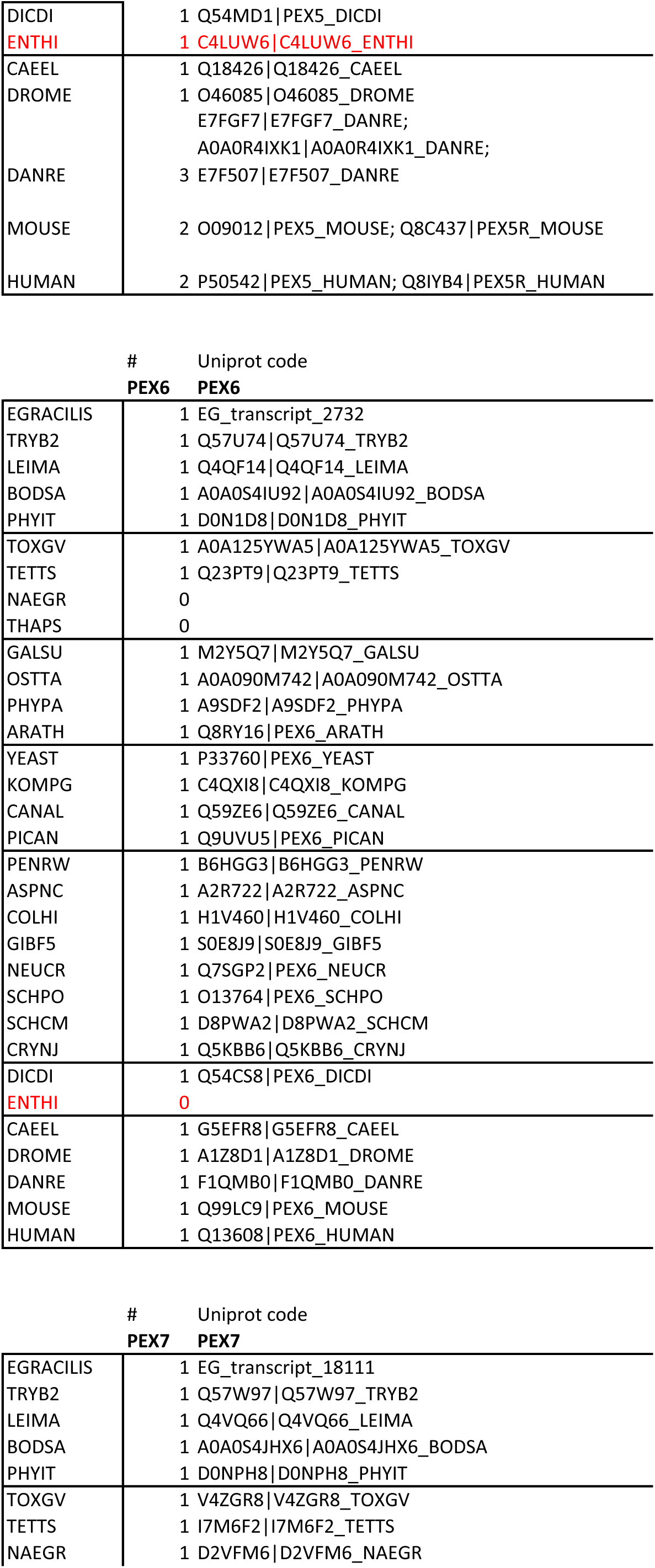
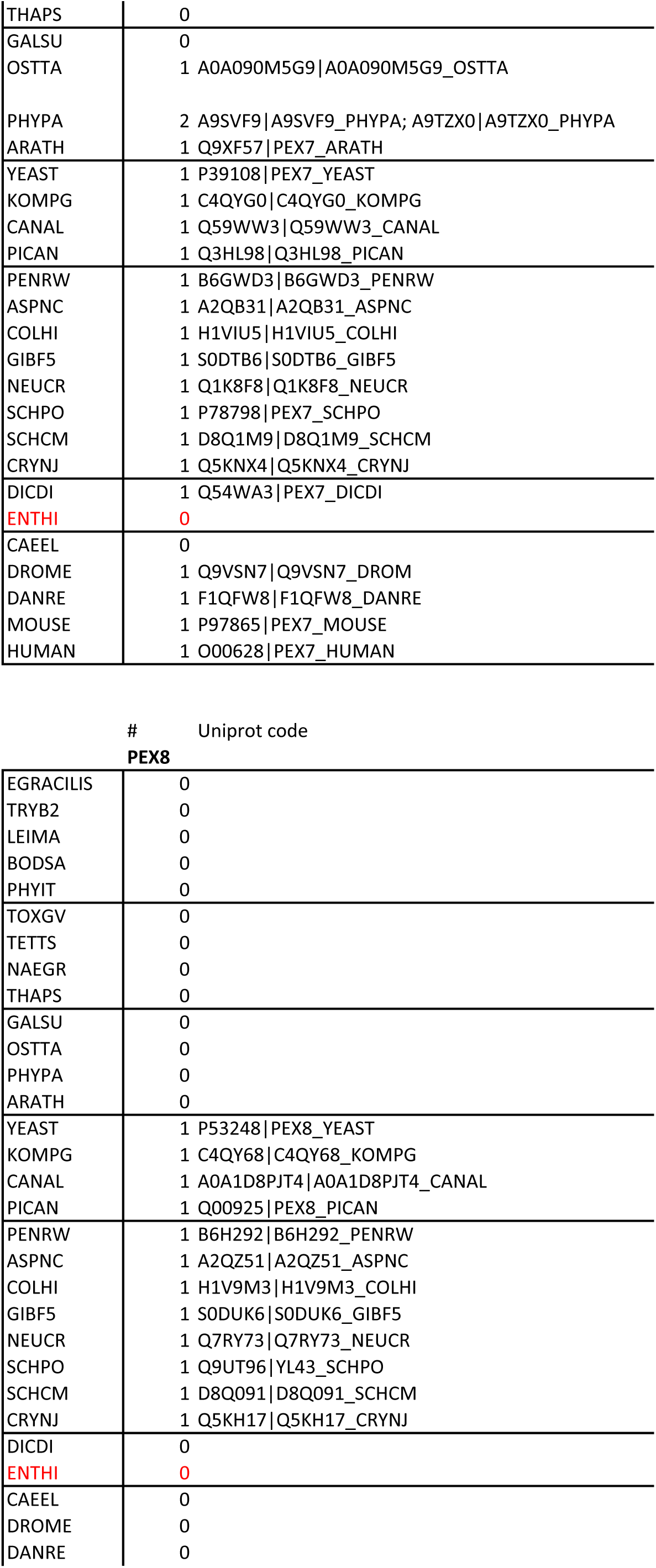
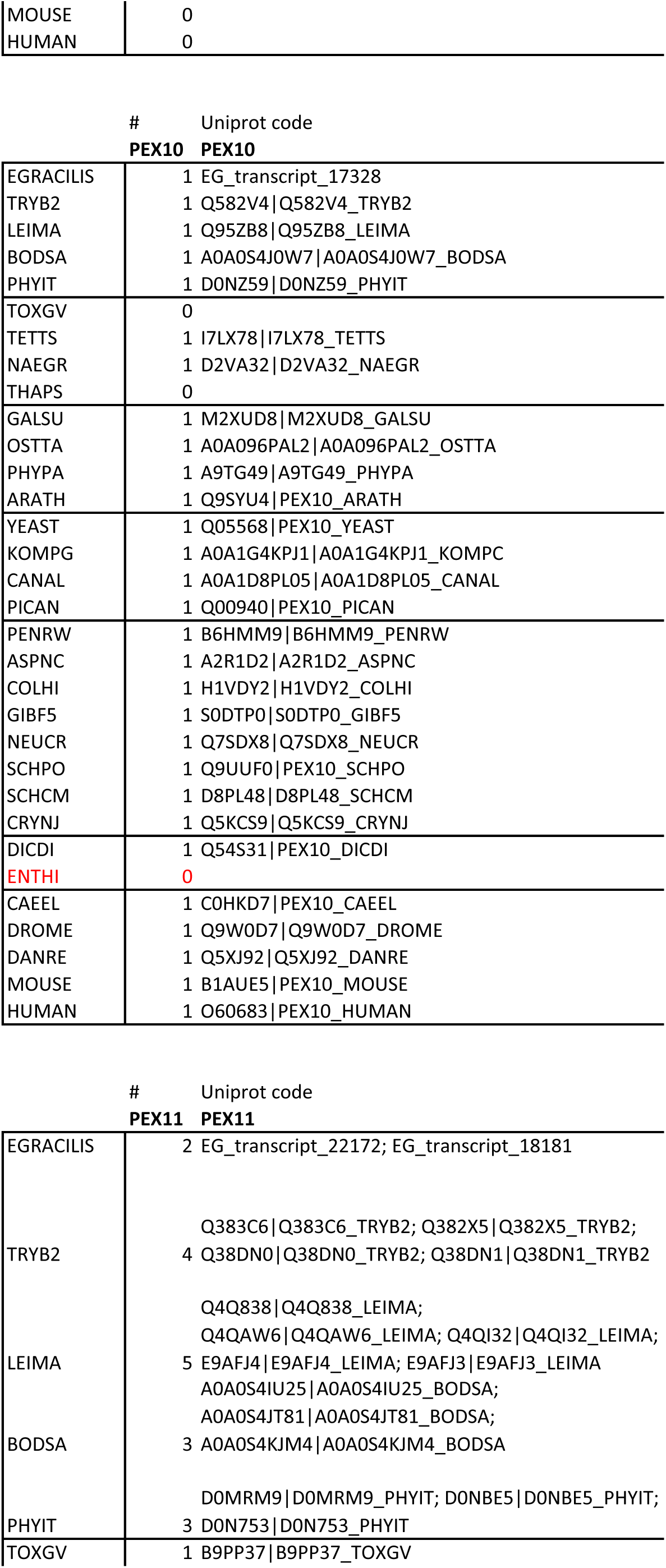
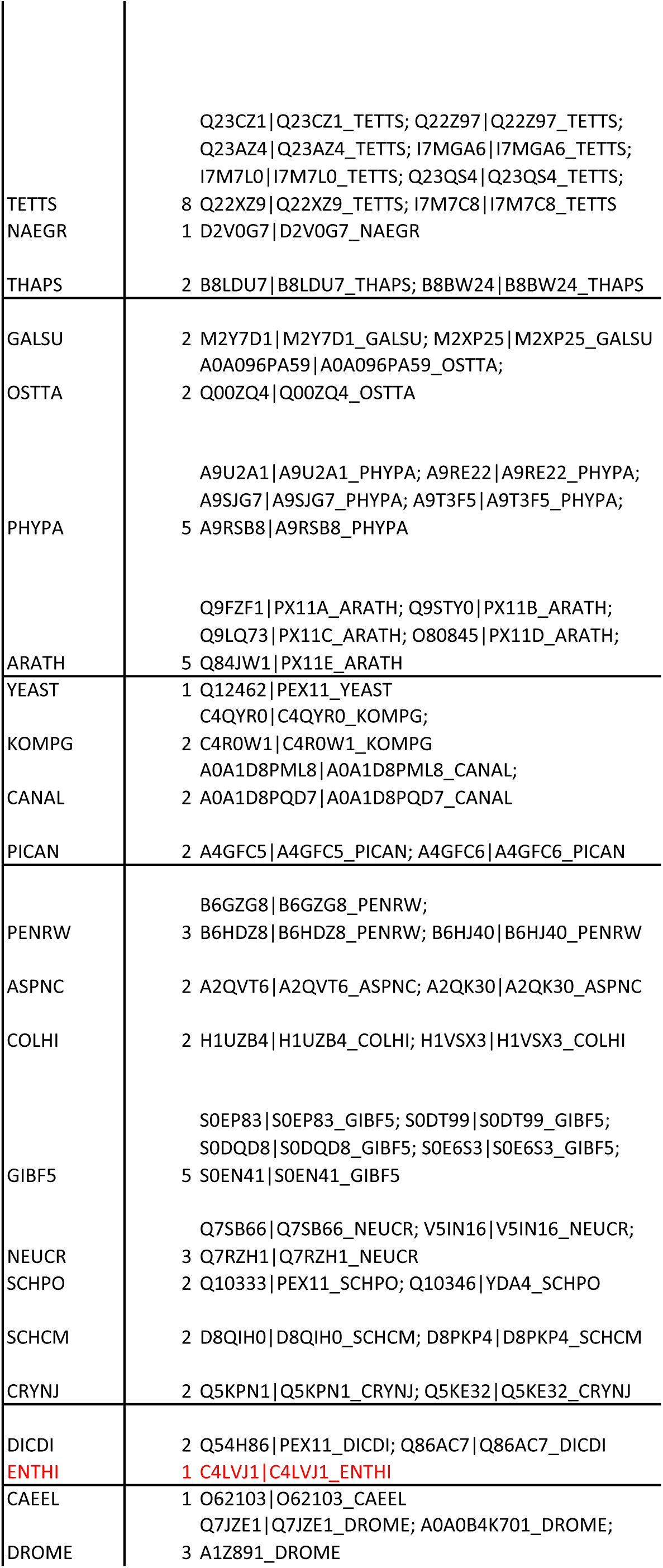
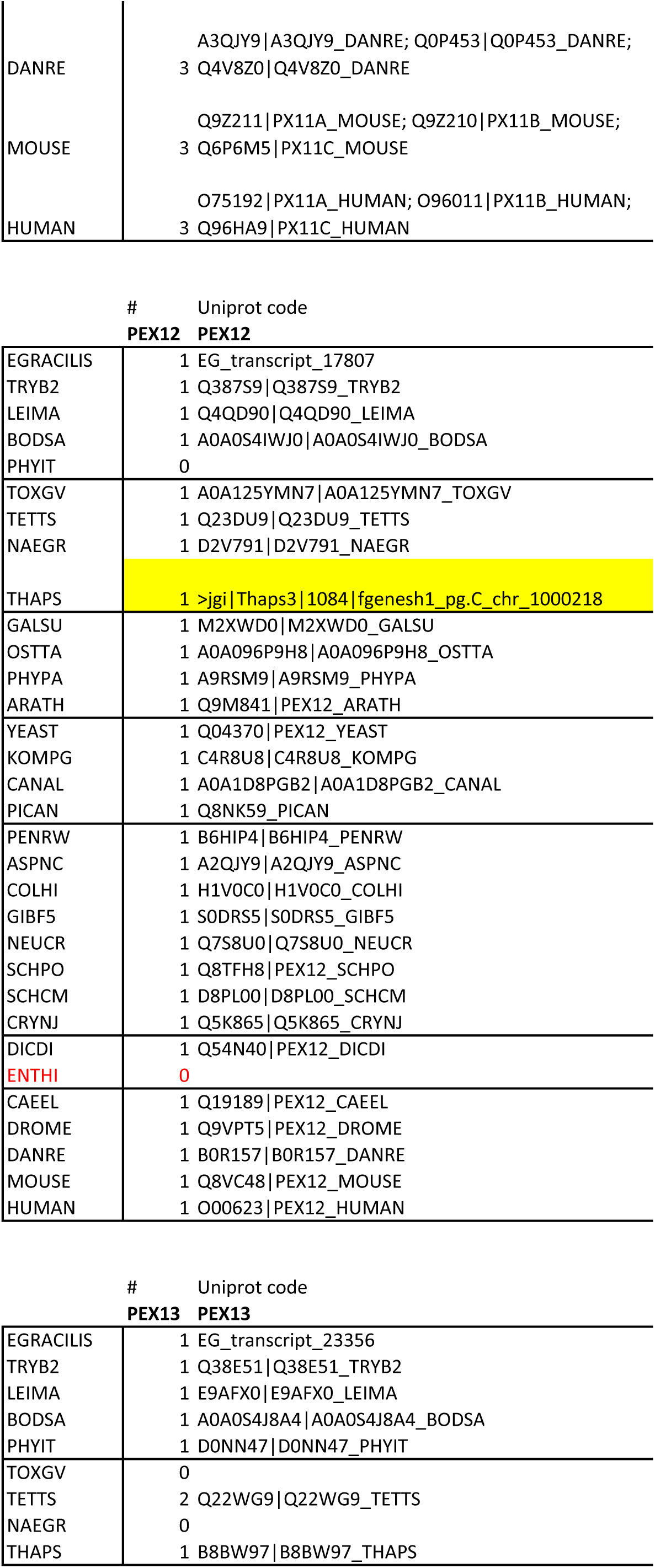
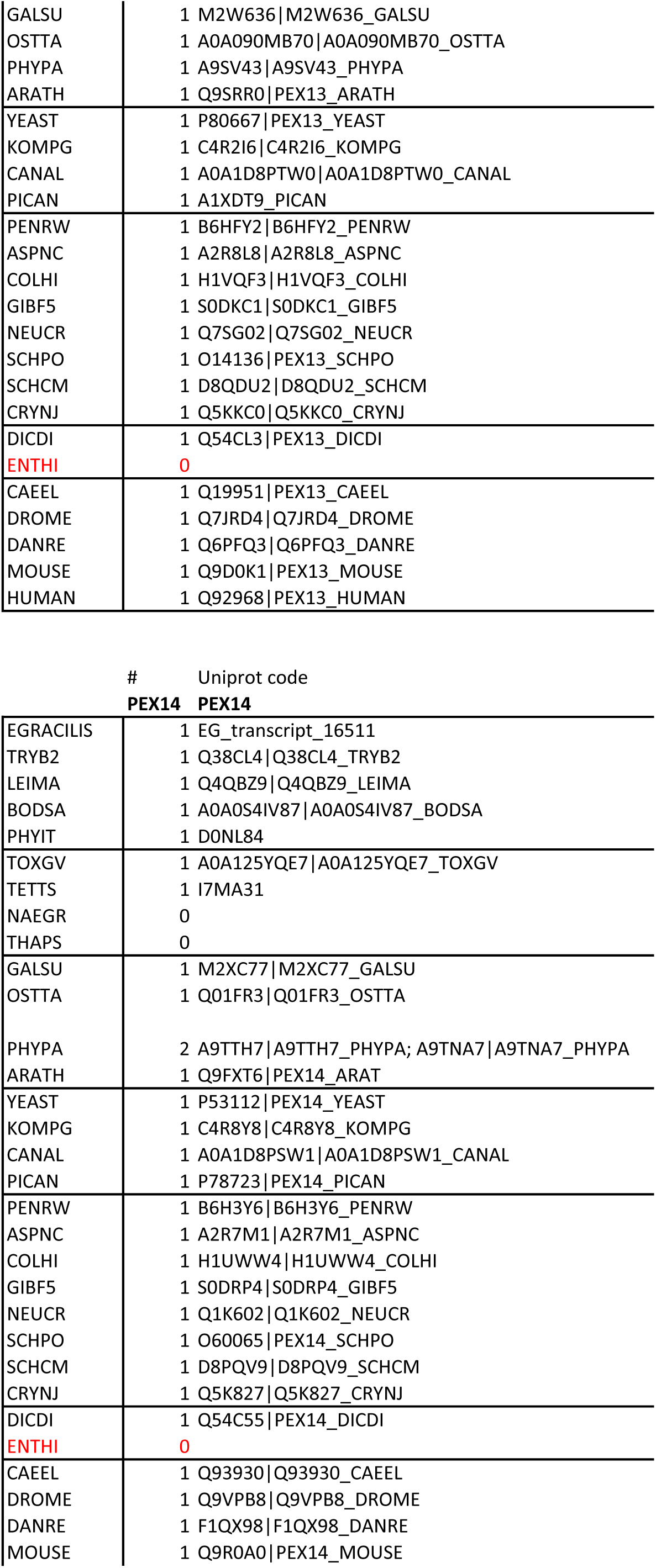
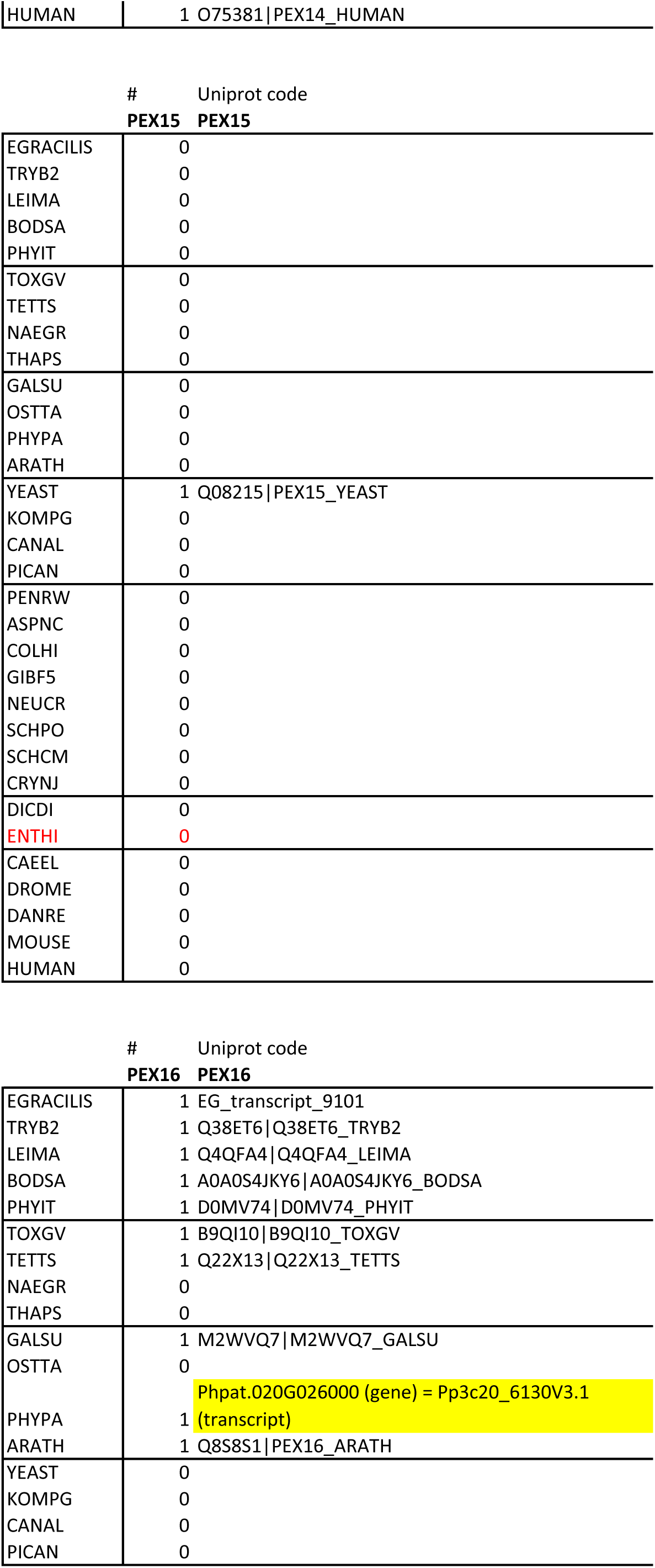
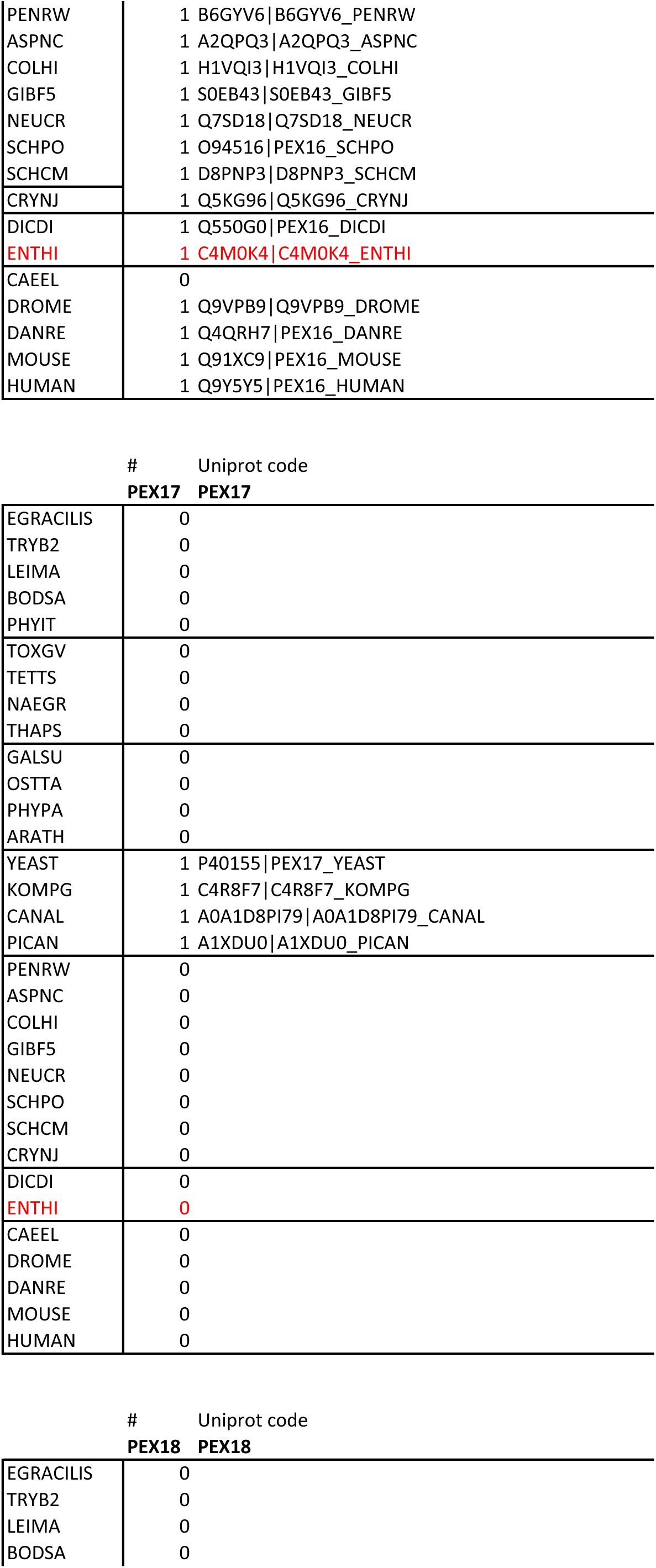
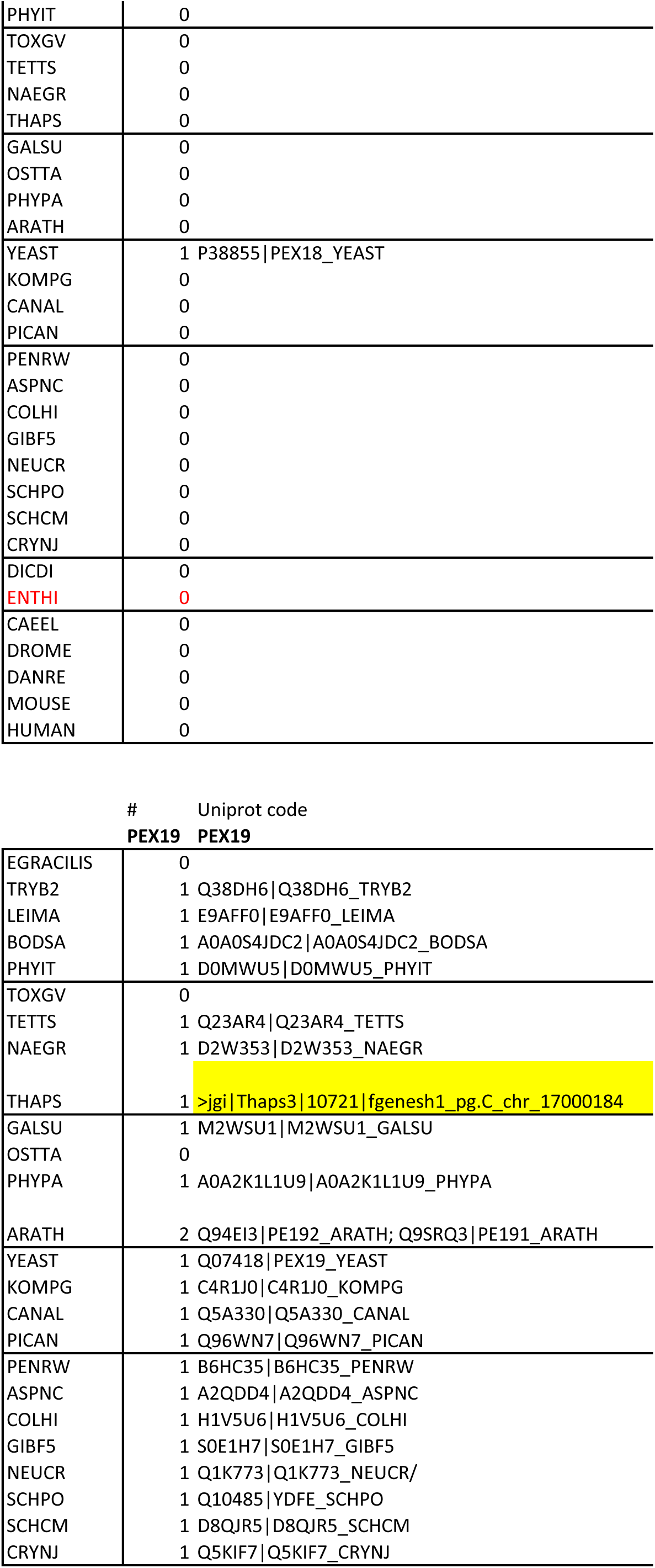
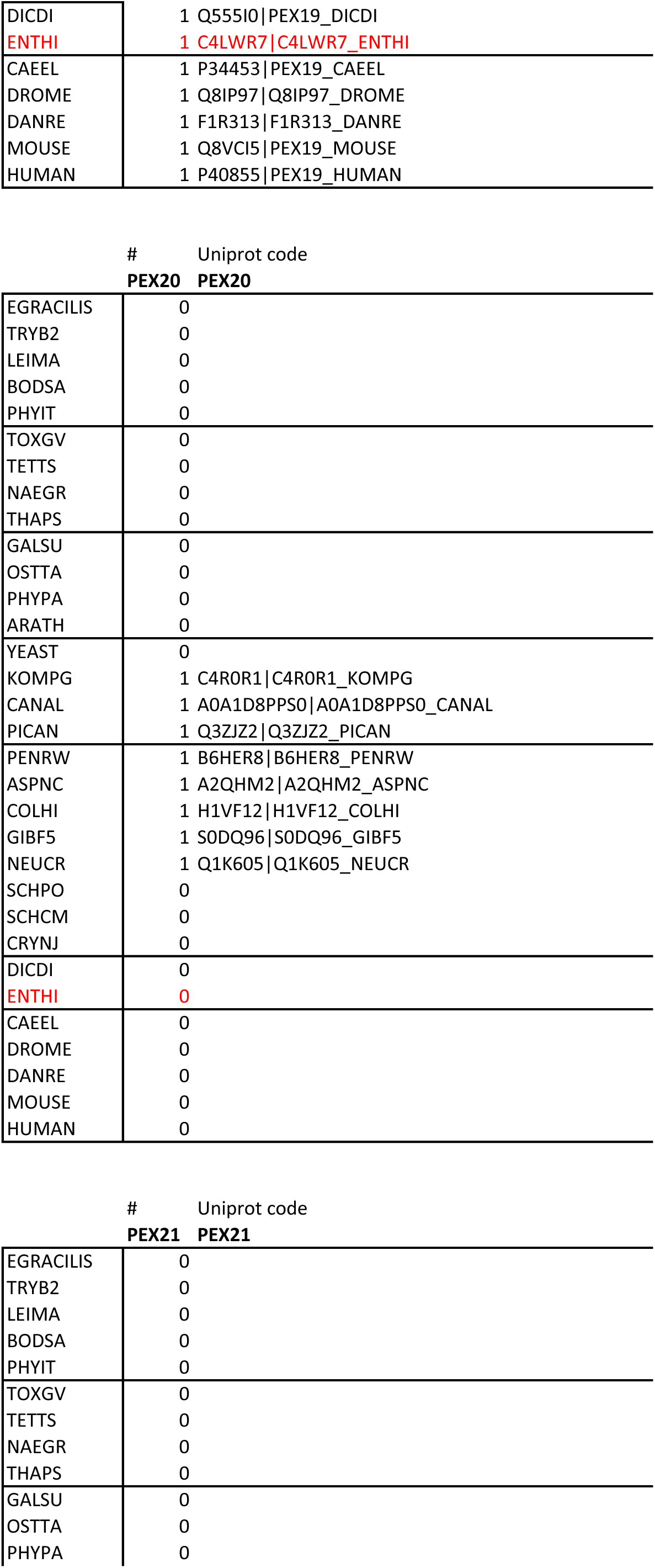
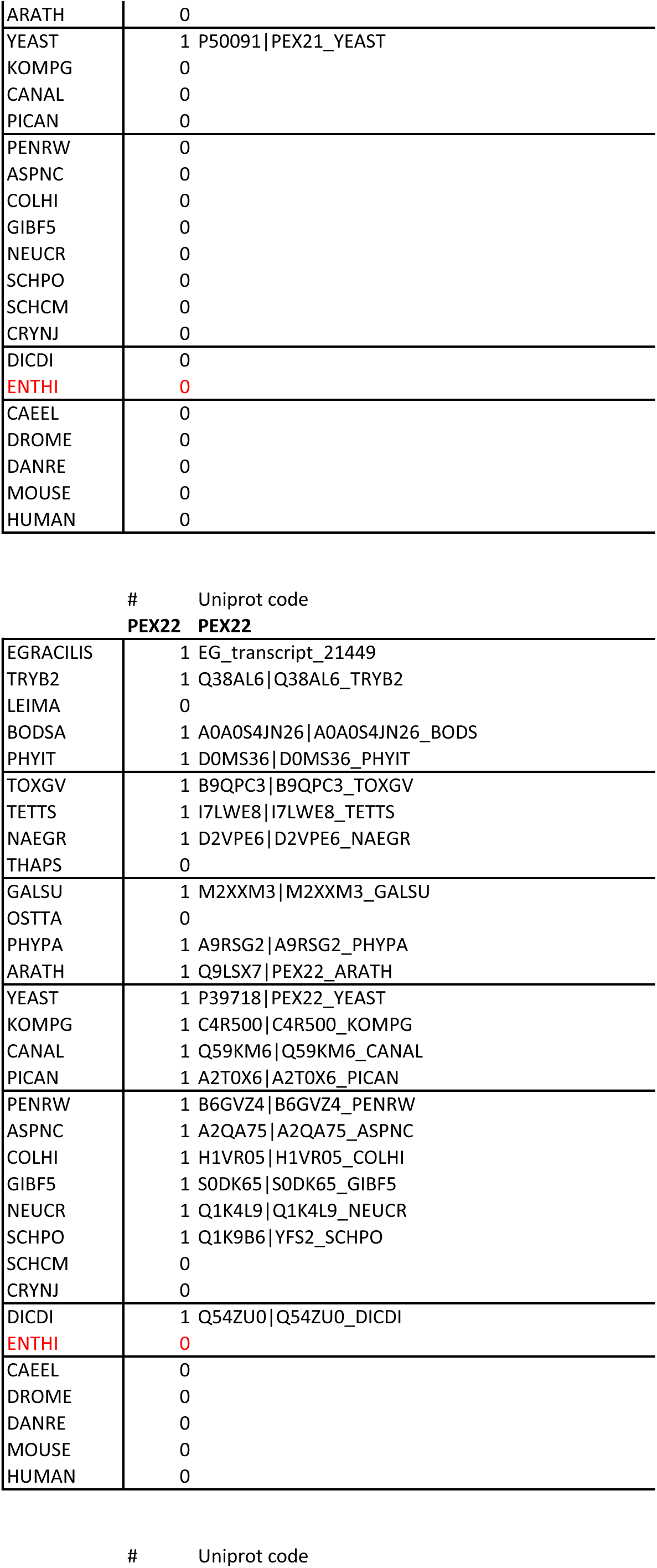
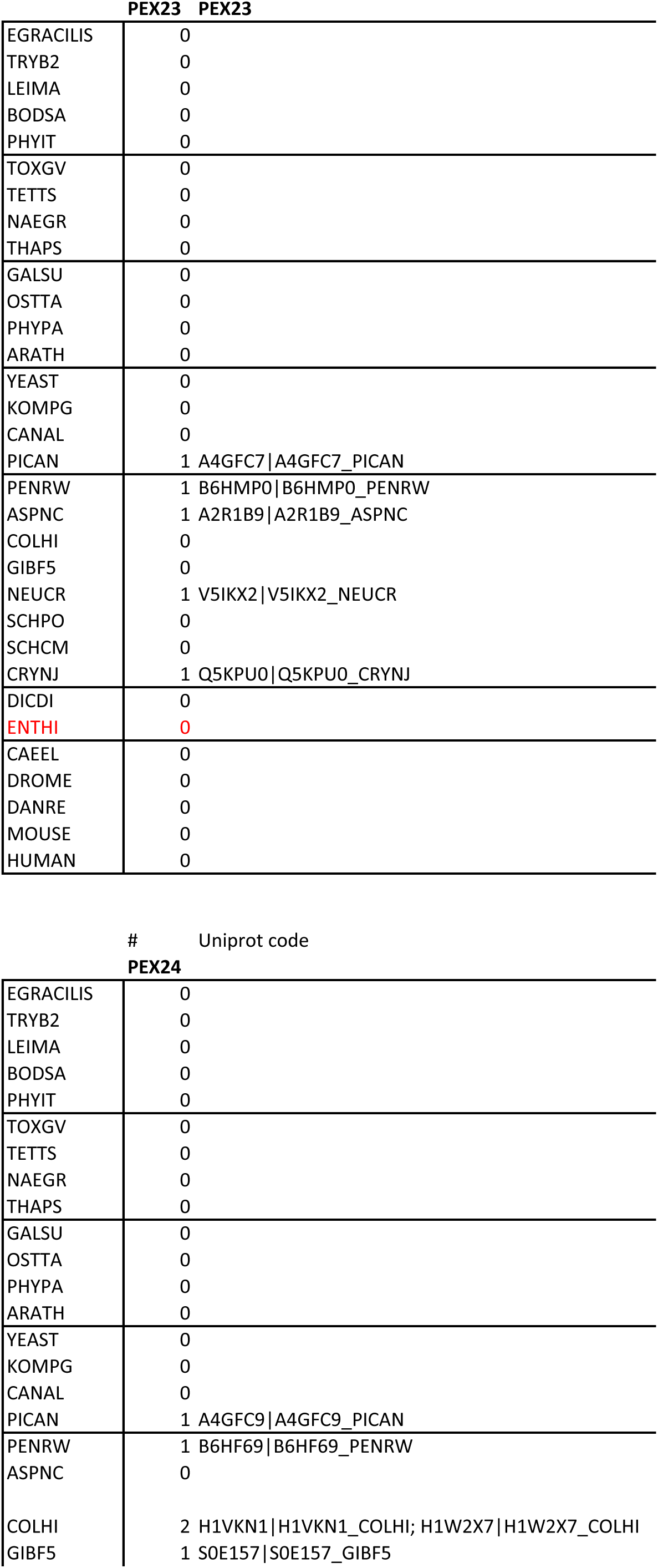
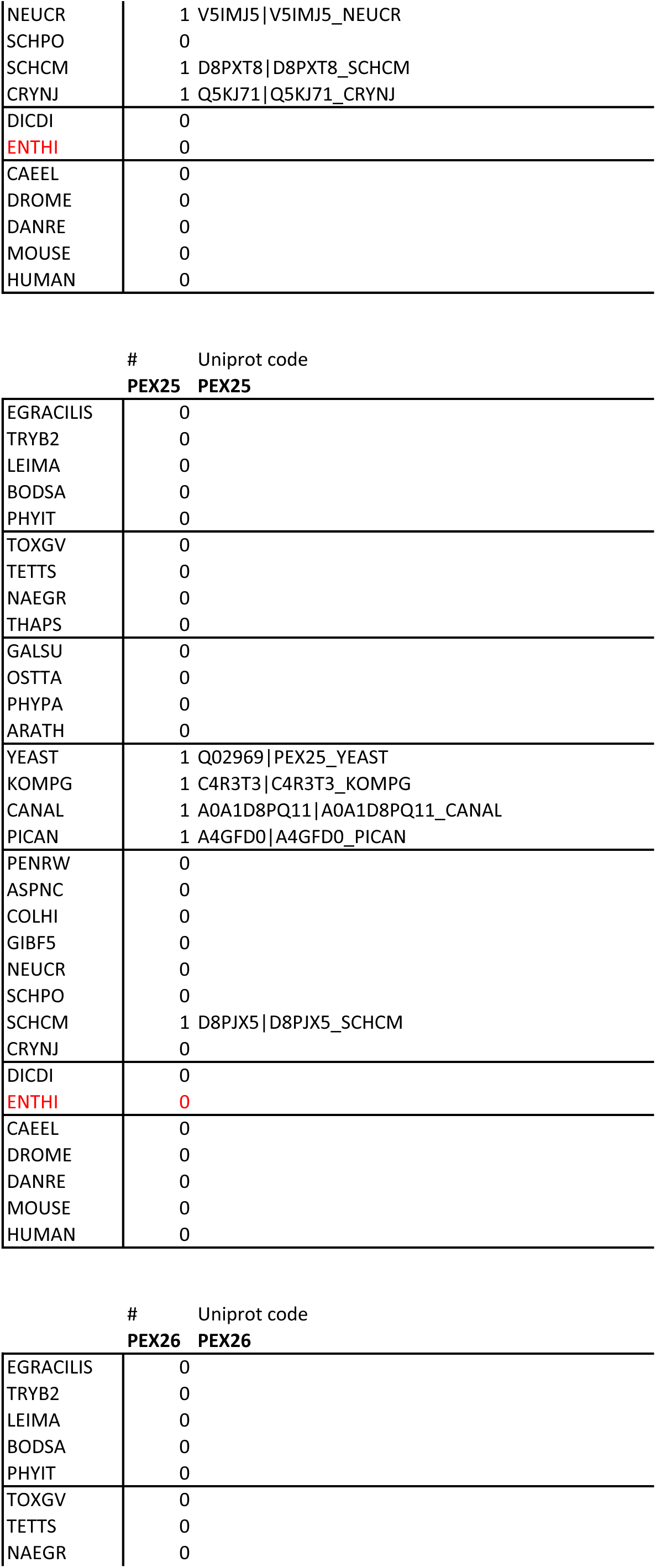
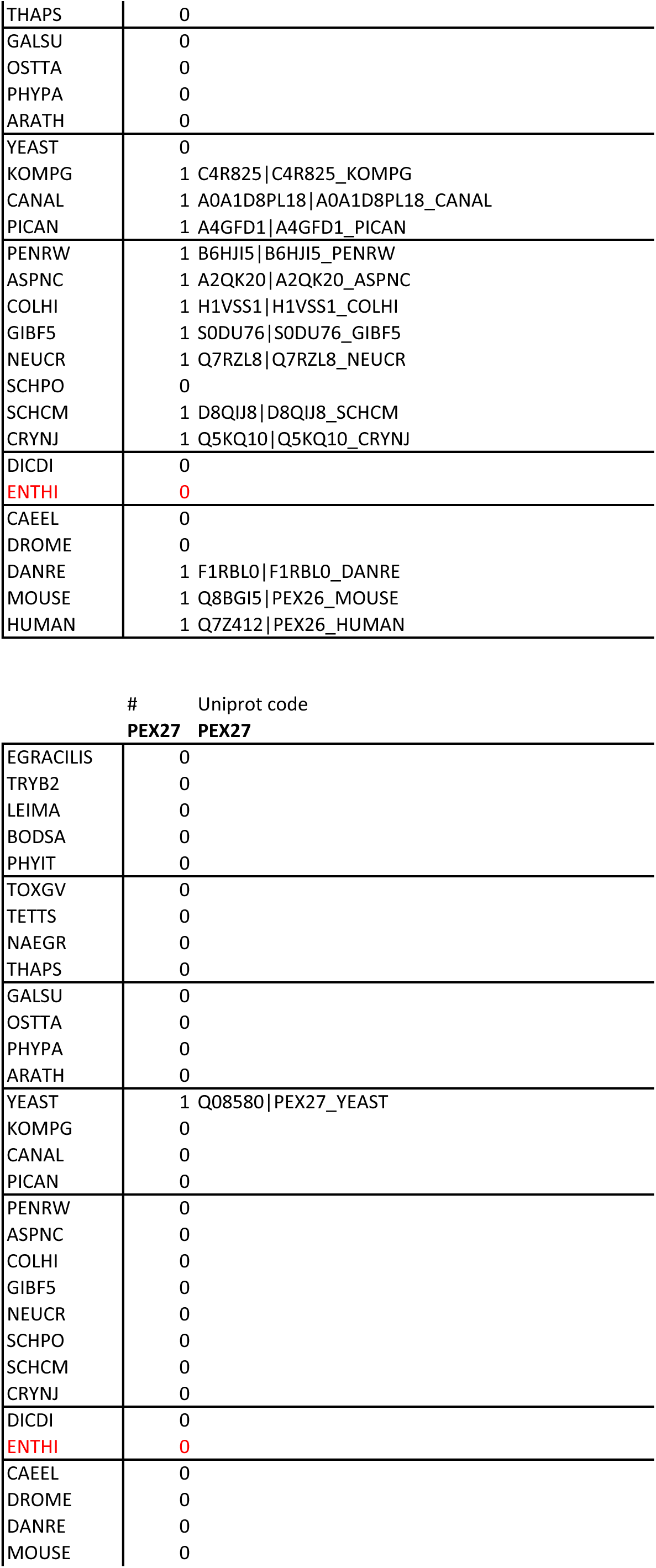
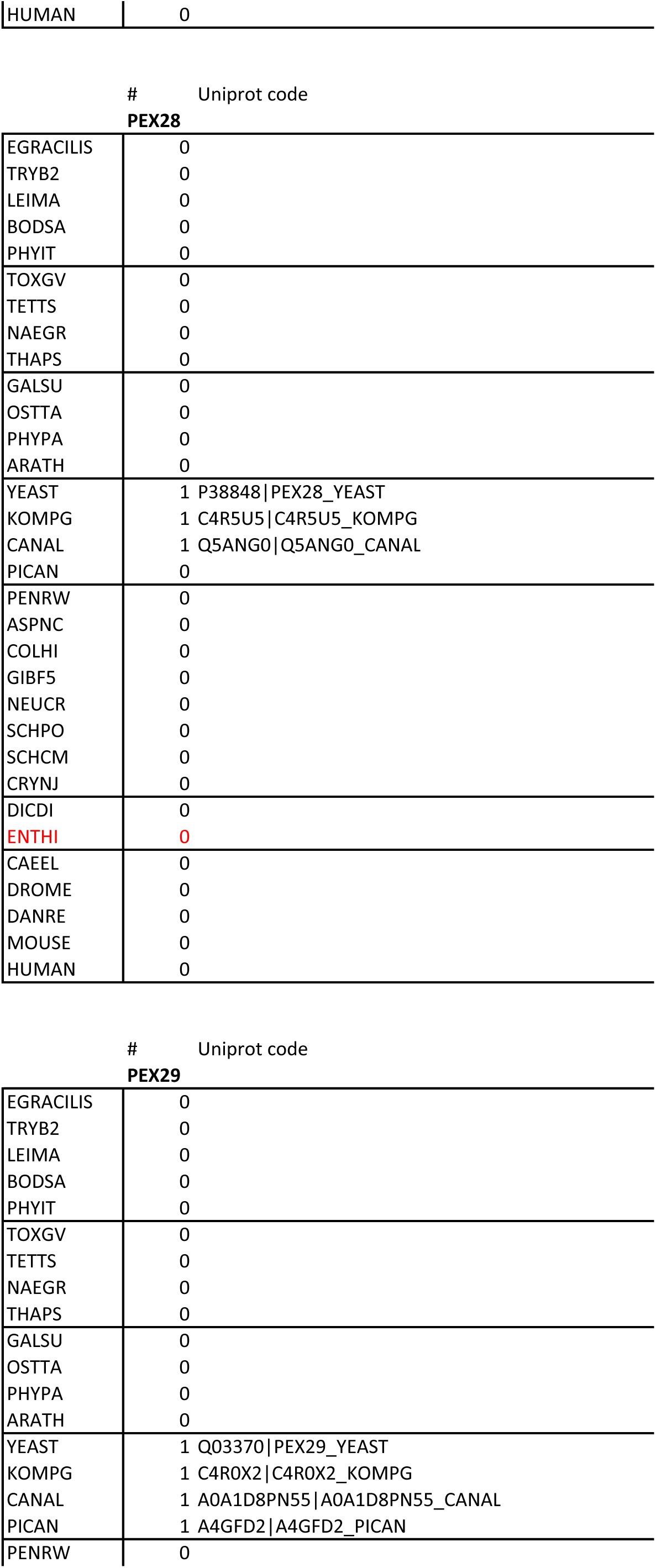
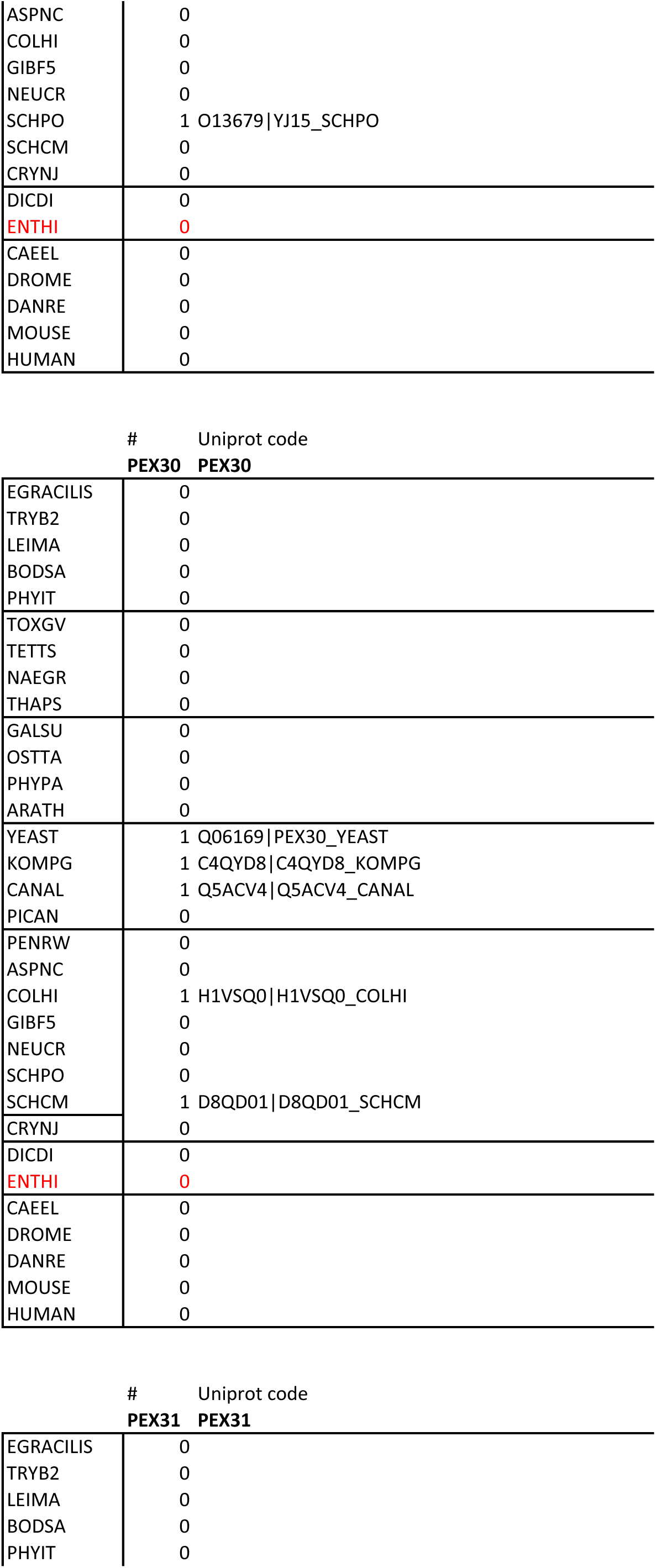
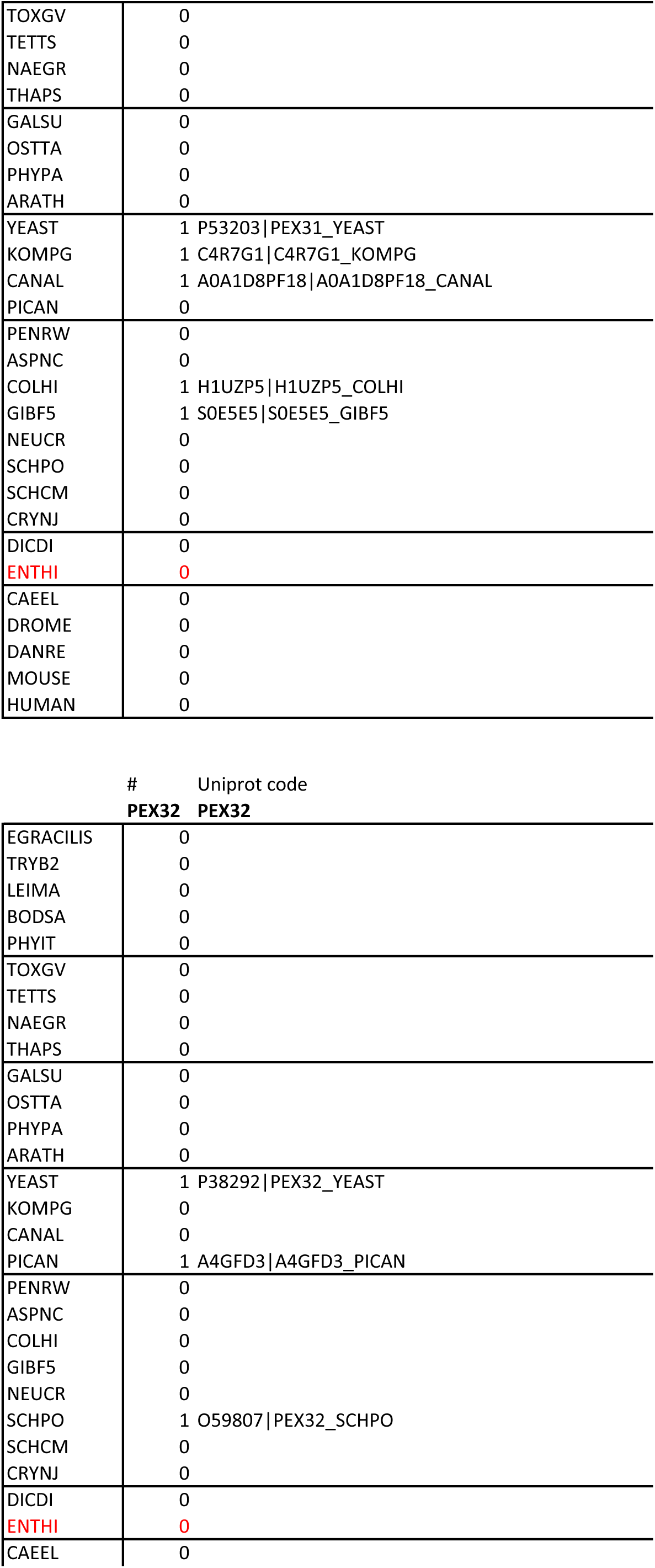
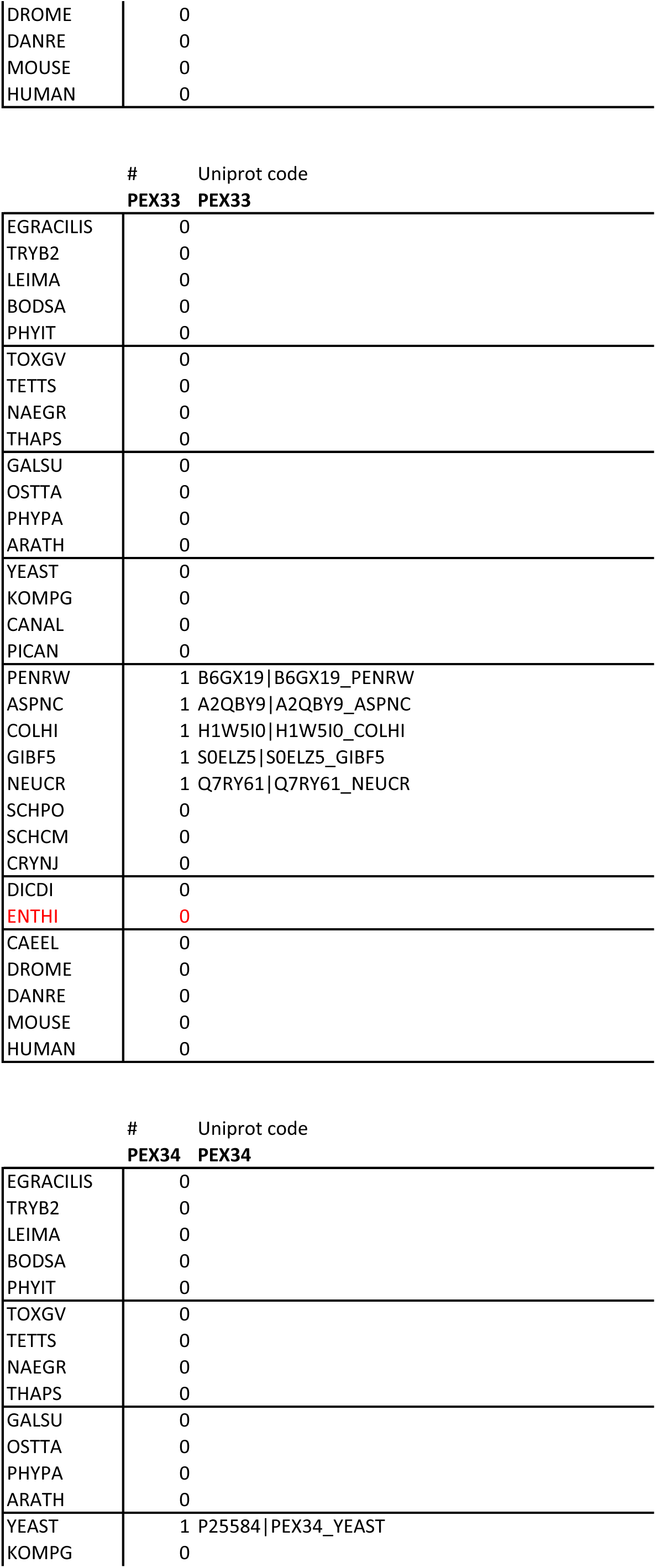
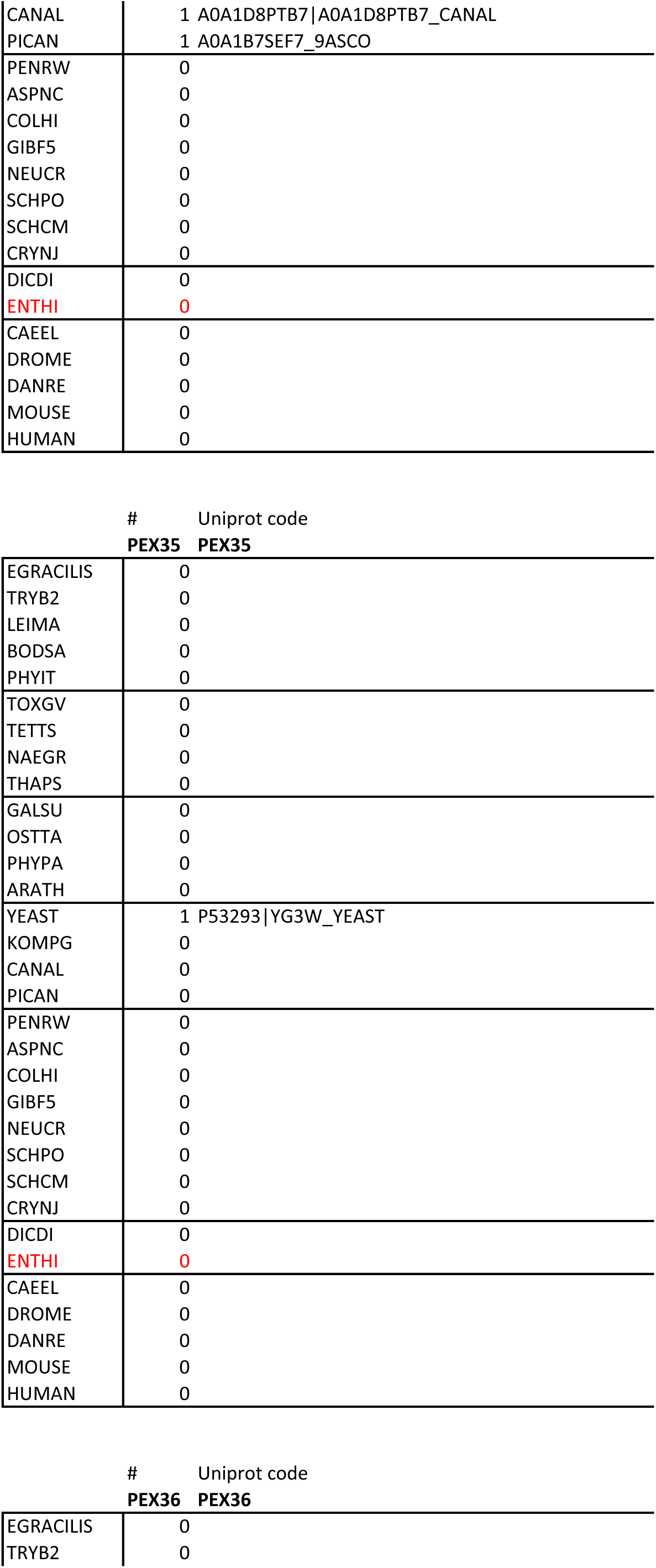
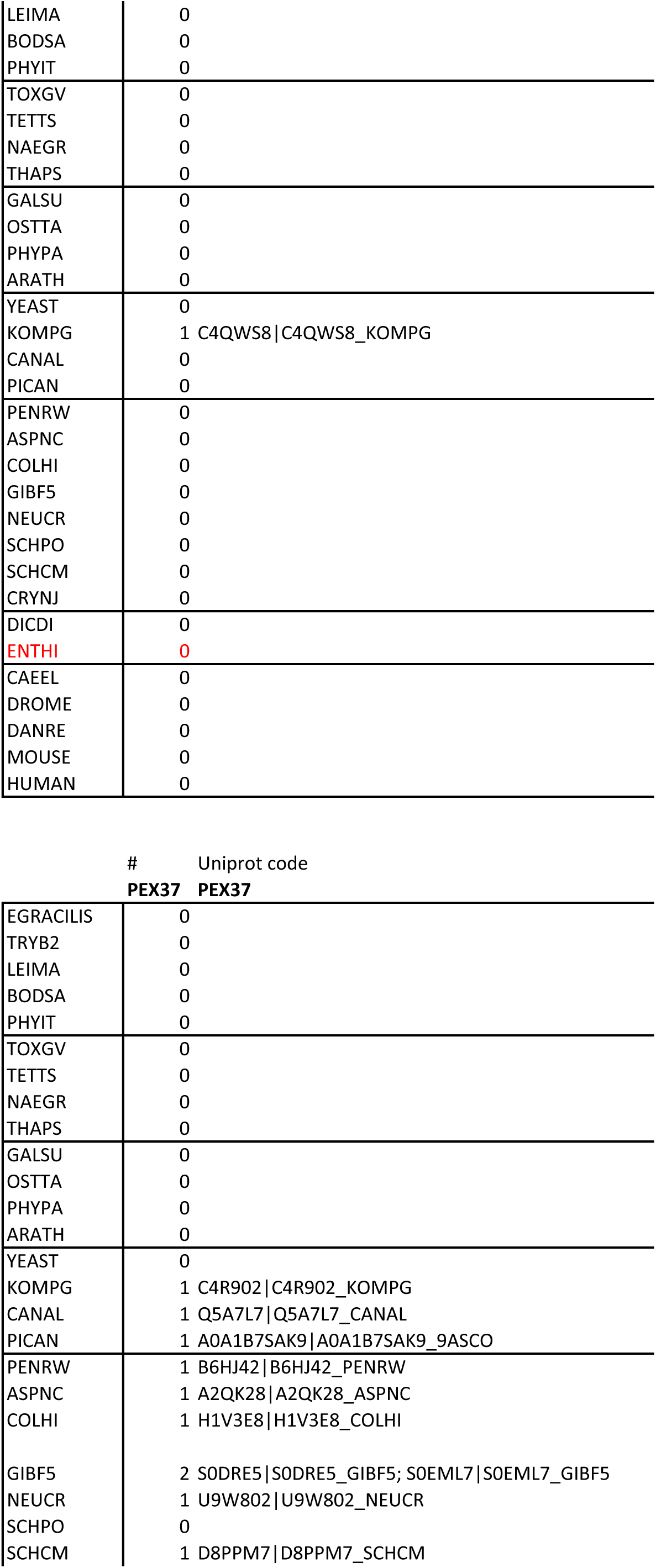
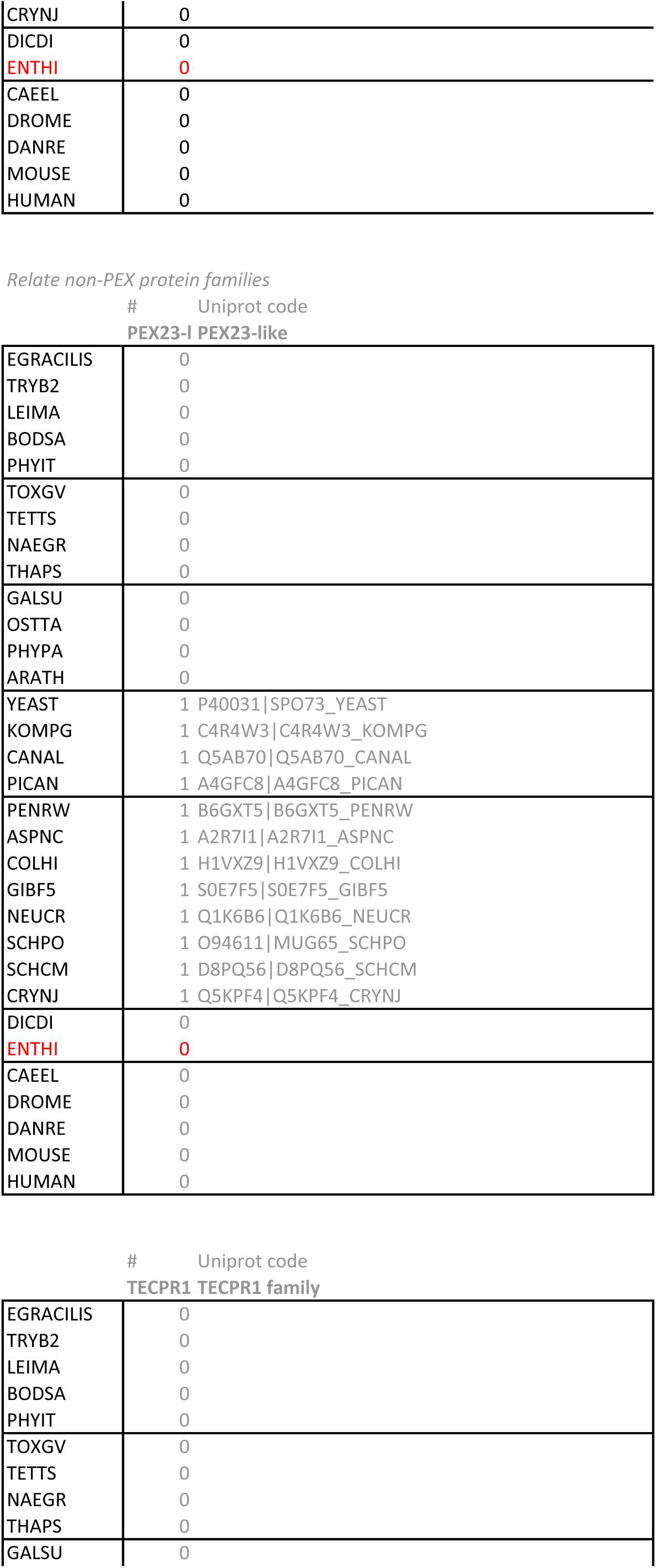
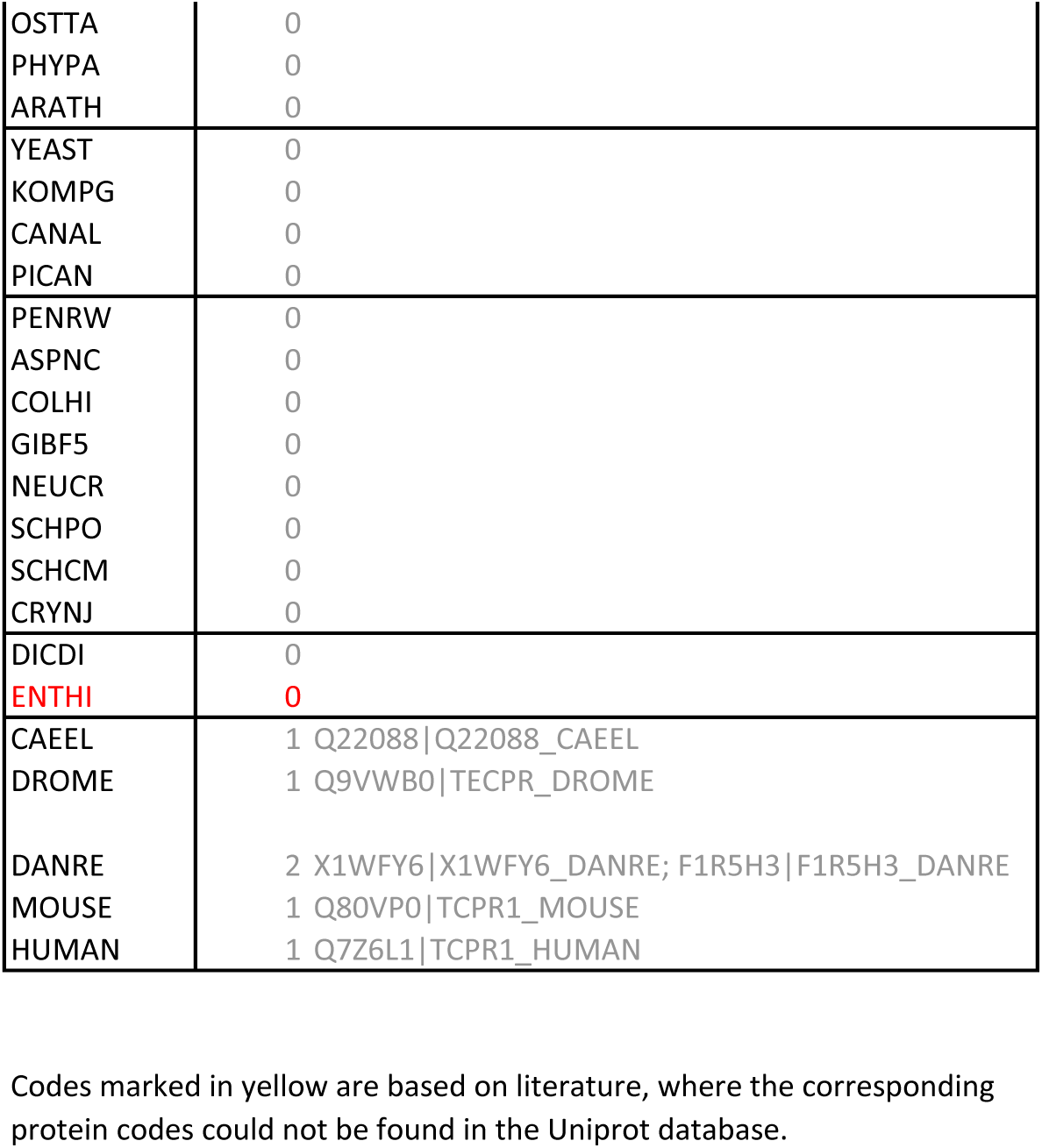
Protein codes for all PEX orthologs (Excel file)

